# Extensive genomic diversity in *Desulfovibrio* species reveals species-specific functional traits associated with disease

**DOI:** 10.64898/2026.03.24.713149

**Authors:** Tingting Zheng, Isabel Keidel, Hélène Omer, Akshat Joshipura, Aurélie Cenier, Ezgi Atay, Yi Han Tan, Daniela Wetzel, Ranko Gacesa, Shijie Zhao, Claudia Mengoni, Shen Jin, Ngoc Anh Khoa Co, Carsten Peters, Nicola Segata, Ruth E. Ley, Monica Yabal, Rinse K Weersma, Dirk Haller, Melanie Schirmer

## Abstract

*Desulfovibrio* spp. are associated with inflammatory diseases and human health, yet limited representative genomes and isolates hinder our understanding of their role in disease. Here, we assembled a comprehensive database of 2,658 *Desulfovibrio* genomes across 90 diseases and 32 countries, including 24 human isolates. Genomic analyses showed extensive species diversity and revealed disease-associated functional traits, including flagellin and virulence genes (i.e. ureases). Flagellin-mediated Toll-like receptor 5 activation was species-specific and *D. desulfuricans* flagellin downregulated TGF-beta signalling in murine small intestinal organoids, suggesting impaired immune tolerance. Additionally, we investigated genomic capacity for hydrogen sulfide (H_2_S) production, a main *Desulfovibrio* metabolite. While health- and disease-associated *Desulfovibrio* spp. mainly encoded dissimilatory sulfate reduction, tetrathionate metabolism-encoding bacteria were exclusively detected in inflammatory bowel diseases, including *Proteus mirabilis* and *Morganella morganii*. Overall, our study provides a comprehensive genomic *Desulfovibrio* resource and identifies new links associating strain variation, functional traits and H_2_S-production with inflammatory diseases.

## Introduction

*Desulfovibrio* are the main sulfate-reducing bacteria in the human gut and increases in their abundance are associated with several diseases, including inflammatory bowel diseases (IBD), colorectal cancer (CRC), systemic sclerosis, Parkinson’s disease and irritable bowel syndrome [1–7]. *Desulfovibrio* spp. promote colonic inflammation in animal models [1, 8, 9] and their relative abundance can discriminate between inflammatory status in mice colonized with patient-derived Crohn’s disease (CD) microbiota [1]. However, several studies have also highlighted positive associations with human health [10–12]. For instance, *Desulfovibrio* abundance was increased in healthy compared to overweight and obese Chinese adults [10] and Swedish preschool children [13]. Furthermore, their abundance was decreased in mucosal biopsies of inflamed tissue from ulcerative colitis (UC) patients compared to non-IBD controls [11]. Collectively, these contradictory findings highlight the need to define the functional role of *Desulfovibrio* in human health and disease.

Detrimental functional traits of *Desulfovibrio* have thus far been mainly studied in the context of hydrogen sulfide (H_2_S), one of its major metabolic products. In the gastrointestinal tract, H_2_S increases cell respiration and is an energy substrate for the mitochondria facilitating detoxification and energy recovery from luminal sulfide [14]. However, excessive H_2_S is cytotoxic, damaging the epithelial barrier and triggering inflammation and colorectal cancer [15]. Bacteria produce H_2_S by metabolizing organic or inorganic sulfur compounds in the human gut. Organic compounds are degraded to release sulfur, which is then converted to H_2_S, while inorganic sulfur compounds can be utilized as terminal electron acceptors to generate energy and H_2_S. Taurine, for example, is an anaerobically respired organic diet-derived sulfur compound [16, 17]. Furthermore, sulfate-reducing gut bacteria, including *Desulfovibrio,* reduce inorganic sulfur compounds, such as sulfite, sulfate, thiosulfate or tetrathionate, to H_2_S [18]. Moreover, H_2_S detoxification proteins, such as sulfide dioxygenase and thiosulfate sulfurtransferase, are downregulated in CD patients [19] and *Desulfovibrio* and dissimilatory sulfate reduction have been linked to high H_2_S concentrations in the stool of colitis-patients [20]. While many studies have focused on dissimilatory sulfate reducers in the human gut, assimilatory sulfate reduction may also play a role in disease. For instance, oxygen-insensitive assimilatory sulfate reduction is more active in fecal samples of CD patients [21]. Bacteria that can employ assimilatory sulfate reduction to produce sulfide include *Escherichia coli* [22], *Salmonella enterica* [23] and *Bacillus subtilis* [24], which are all highly adaptive and persistent in the human gut environment. Furthermore, cysteine degradation leads to H_2_S production and a wide range of putative cysteine-degrading gut bacteria were recently identified [25]. Another important H_2_S source is tetrathionate respiration, where tetrathionate is first reduced to thiosulfate through the *ttr* operon (*ttrBCA*) and then to H_2_S [26]. Interestingly, tetrathionate reduction provides a growth advantage to *Salmonella typhimurium* in mice [27]. Despite these recent advances, the identification of H_2_S contributors in the human gut remains underexplored.

Other studies investigating functional traits have focused on individual *Desulfovibrio* strains. For example, *Desulfovibrio* QI0027 is able to fix nitrogen, which may play a role in the availability of nitrogen in the gut [28] and *Desulfovibrio desulfuricans* lipopolysaccharides induce interleukin(IL)-6 and IL-8 from human endothelial cells [29]. Cross-feeding between *Desulfovibrio piger* and *Faecalibacterium prausnitzii* revealed that acetate production by *D. piger* enhances *F. prausnitzii* growth and butyrate production [30]. However, as *Desulfovibrio* spp. remain difficult to isolate, these findings rely on individual *Desulfovibrio* strains, while a comprehensive genomic analysis is still lacking.

Only 280 *Desulfovibrio* genomes are present in the NCBI RefSeq database (February 2026). Furthermore, the family Desulfovibrionaceae was recently reclassified from Deltaproteobacteria to the new class Desulfovibrionia [31], emphasising the need for careful investigation of existing taxonomic annotations in databases. Reconstructing genomes from metagenomic sequencing data is an important approach to study genomic diversity of microbial species, especially for organisms that remain difficult to culture [32, 33]. For example, a tree of life was constructed based on metagenome-assembled genomes (MAGs) from a variety of different environments, which revealed a large number of lineages exclusively recovered as MAGs [32]. Furthermore, this approach also substantially expanded our knowledge of species, such as *Segatella copri* [34], *Akkermansia muciniphila* [35], *Eubacterium rectale* [36] and *Bifidobacterium* spp. [37], associating genomic diversity within these species to ecological function, host lifestyle, and health.

In this study, we constructed a comprehensive database of 2,658 *Desulfovibrio* genomes and assembled a collection of 24 human isolates from 8 major *Desulfovibrio* species. For this, metagenome-assembled genomes (MAGs, n=2,499) were recovered from >100.000 cohort samples, in addition to 442 genomes from major public genome databases. Overall, our database represents *Desulfovibrio* genomes recovered from 174 studies across 90 disease phenotypes and 32 countries. We identified 37 *Desulfovibrio* species, including 3 potential new species, which showed high gene diversity in cell motility, virulence, and antimicrobial resistance. Comparative genomic analyses identified IBD- and CRC-associated functional traits, including urease and flagellar genes. Flagellated *D. desulfuricans* and *D. legallii* showed strong activation of Toll-like receptor 5 (TLR5), a key player in innate immunity. Recombinantly expressed flagellin proteins further confirmed flagellin-specific TLR5 activation. Interestingly, flagellin-treated murine small intestinal organoids showed suppression of the TGF-beta signalling pathway by *D. desulfuricans* flagellin, which may lead to disruptions of both innate and adaptive immune tolerance. Geographic differences in gene content of *Desulfovibrio sp900319575* were also observed, including cell motility, virulence and genome plasticity genes. Furthermore, we investigated genomic gut microbial H_2_S production capacity. This revealed that dissimilatory sulfate reduction was well conserved across *Desulfovibrio* genomes, but with no disease-association. However, tetrathionate metabolism genes were increased in abundance and transcriptionally active in CD patients. We identified 9 genera encoding the complete tetrathionate metabolism pathway, including disease-associated *Proteus mirabilis* and *Morganella morganii*, offering new insights into their potential role in disease. In summary, our study reveals extensive genomic diversity and facilitates functional genomic analyses of *Desulfovibrio* to uncover their role in human health and disease.

## Results

### Constructing a comprehensive *Desulfovibrio* database of 2,658 genomes

For a comprehensive characterization of *Desulfovibrio* spp., we applied a MAG recovery approach. First, we analysed 11,017 samples from 10 diverse human metagenomic cohort studies, resulting in 1,921 new *Desulfovibrio* high-quality MAGs (**Supplementary Table 1**+**2**). These cohorts span different diseases (including IBD and CRC), ages (young, elderly and centenarians) and geographical areas (i.e. Southern China, Japan, Nepal, Tanzania, the Netherlands, the US, Germany, France and Israel). The majority of the recovered MAGs (1726/1921; 89.85%) came from two Dutch cohorts. High *Desulfovibrio* prevalence was observed in both Dutch studies (LifeLineNext [38] mother & infant study, prevalence in adults = 26.67%; Dutch microbiome cohort [39], prevalence = 20.86%), while several Chinese populations showed a high degree of variation in prevalence (max prevalence: 60%, min prevalence: 5%, **Supplementary Table 3**). Interestingly, sequencing depth was not a major driver of *Desulfovibrio* spp. detection (**Supplementary Fig. 1+2**). In addition, 578 *Desulfovibrio* genomes were recovered from MetaRefSGB[40] (**Supplementary Table 1**), a resource that includes a collection of MAGs from >100k metagenomes spanning different body sites, ages, countries and lifestyles. Furthermore, we identified 442 *Desulfovibrio* genomes in publicly available databases, including NCBI RefSeq, the Unified Human Gastrointestinal Genome (UHGG), HRGM, IMG/M and MGnify. Due to recent re-classification efforts, we re-annotated all of these genomes using GTDB-tk annotations [41]. This led to the re-assignment of 31 genomes to other genera within Desulfovibrionaceae (**Supplementary Table 4**), which were subsequently excluded from our analysis. Overall, this genome database represents *Desulfovibrio* diversity recovered from human, animal, and environmental samples (**Fig. 1a**).

**Fig. 1.**
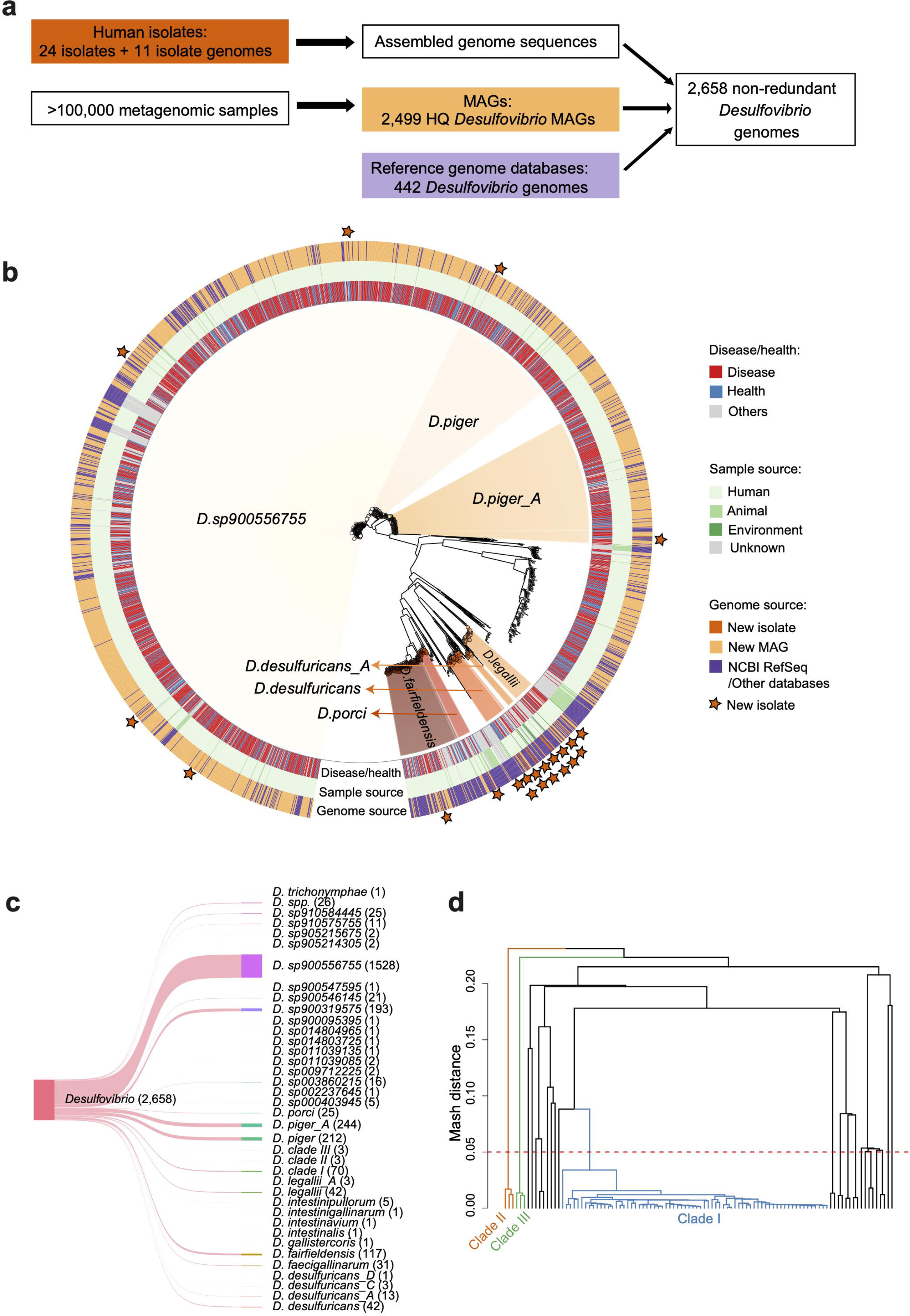
Genomic analysis of 2,658 *Desulfovibrio* genomes. **a.** We reconstructed a *Desulfovibrio* database consisting of 1,921 high-quality metagenome-assembled genomes (MAGs) identified from the analysis of 11,017 metagenomic samples from 10 human cohort studies with different health conditions (e.g. healthy individuals, inflammatory bowel diseases [IBD], colorectal cancer [CRC], and other diseases) and geographical areas, including Southern China, Japan, Nepal, Tanzania, the Netherlands, the US, Germany, France and Israel. In addition, 578 *Desulfovibrio* genomes were retrieved from MetaRefSGB, containing MAGS from >100k metagenomes [40], and major public genome databases. Furthermore, we isolated and obtained 24 *Desulfovibrio* strains from human cohort samples and public culture collections, which were subsequently sequenced. Additionally, 11 human isolate genomes were downloaded from NCBI RefSeq. All available genomes were merged and duplicate entries removed, resulting in 2,658 *Desulfovibrio* genomes. HQ MAGs - high-quality metagenomic assembled genomes. **b.** Phylogenetic tree of all *Desulfovibrio* genomes based on mash distance. Rings indicate host phenotype (diseased *vs.* healthy, “others” refers to genomes from animals, natural environments or unknown origin), sample source (human *vs.* animal *vs.* environment) and genome source (newly identified MAG *vs*. newly sequenced isolate or database entry). Stars indicate strains for which isolates are available. Species names follow GTDB taxonomy **c.** Overview of the number of genomes within each species. The species with the most genomes is *D. sp900556755* (proposed name: *Desulfovibrio longus* sp. nov.). followed by *D. piger_A*, *D. piger*, *D. sp900319575*, and *D. fairfieldensis.* **d**. Hierarchical clustering of 86 genomes from uncharacterized *Desulfovibrio* spp. based on mash distance (cutoff <5%). Colors indicate *Desulfovibrio* clades: clade I (blue), clade II (orange), and clade III (green).

As isolates are crucial for any type of functional validation experiments, we next focused on culturing and procuring human *Desulfovibrio* isolates. We were able to isolate 2 strains from IBD and CRC patients and 7 strains from healthy individuals. In addition, 15 clinical isolates were acquired from microbial culture repositories, leading to a total of 24 newly sequenced human isolates representing 8 major *Desulfovibrio* species and one uncharacterized *Desulfovibrio* spp. (**Fig. 1a**, **Supplementary Table 5**).

After merging all available genomes, they were de-replicated (99.9% average nucleotide identity [ANI] cutoff [42, 43]), resulting in a final curated database of 2,658 unique, high-quality *Desulfovibrio* genomes, spanning 37 species (including new candidate species, **Fig. 1b+c**), recovered from 174 metagenomic studies (including 101 human microbiome studies) from 32 countries, representing 90 different diseases and health conditions (**Supplementary Table 1 and 6)**. Most genomes belonged to *Desulfovibrio longus* sp. nov. (i.e. *D. sp900556755* based on GTDB-tk) (n=1,528, 57.49%), which constituted the major species represented in this database. Other main species (containing >50 genomes) were *D. piger_A*, *D. piger*, *D. sp900319575*, and *D. fairfieldensis* (**Fig. 1b+c**). In summary, this comprehensive *Desulfovibrio* database with its host-associated information will serve as the basis for exploring genomic diversity and strain-and gene-specific disease signals, which can then be further investigated using the representative clinical isolates.

### *Desulfovibrio* spp. are found across different human diseases and health conditions

To investigate *Desulfovibrio* species diversity and phylogeny, we calculated pairwise mash distances between all genomes (**Fig. 1b**). While *Desulfovibrio* is often associated with disease [1–4], they were also prevalent in healthy individuals with 686 *Desulfovibrio* genomes recovered from healthy samples versus 1,503 recovered from patient samples. However, no obvious clustering by disease phenotype was observed, indicating that overall genome similarity is insufficient and detailed analyses are required for the identification of disease-specific functional traits. Interestingly, *Desulfovibrio sp910575755* and *Desulfovibrio sp910584445*, two murine species, clustered separately, potentially reflecting host adaptation, while 14 environmental *Desulfovibrio* genomes (originating from soil, sand, sediment, freshwater, wastewater, bioreactor and agricultural waste), were interspersed with genomes from human samples.

Furthermore, our analysis identified 3 potential novel species. Among the 102 *Desulfovibrio* genomes that could not be assigned to any existing species, 76 genomes clustered into 3 clades (mash distance <5% [44–46], **Fig. 1d**). *Desulfovibrio* clade I contained 70 genomes, including 58 MAGs from the two Dutch cohorts, while the other two clades were smaller with 3 MAGs from the Dutch cohorts and 3 MAGs from a Hadza hunter population, respectively. The detection of these novel species highlights that *Desulfovibrio* spp. are highly diverse and that much of this diversity remains to be uncovered.

### *Desulfovibrio* spp. differ in cell motility, ureases, NADH dehydrogenase and oxidoreductase genes

We next identified differential genetic features in different *Desulfovibrio* species. Several clusters of orthologous groups (COGs) emerged, in particular COGs related to cell motility (**Fig. 2a**), suggesting species-specific difference in motility. Especially a flagellin gene (COG1344) was present in all genomes for 28 *Desulfovibrio* species, including *D. fairfieldensis* (117/117), *D. desulfuricans* (42/42), *D. legallii* (42/42) and *Desulfovibrio* clade III (3/3), while completely absent in other species, including *D. piger*, *Desulfovibrio longus* sp. nov., and *Desulfovibrio* clade I and II. This indicates increased motility and potential microbe-host interactions [47] of *D. fairfieldensis, D. desulfuricans*, *D. legallii* and *Desulfovibrio* clade III.

**Fig. 2.**
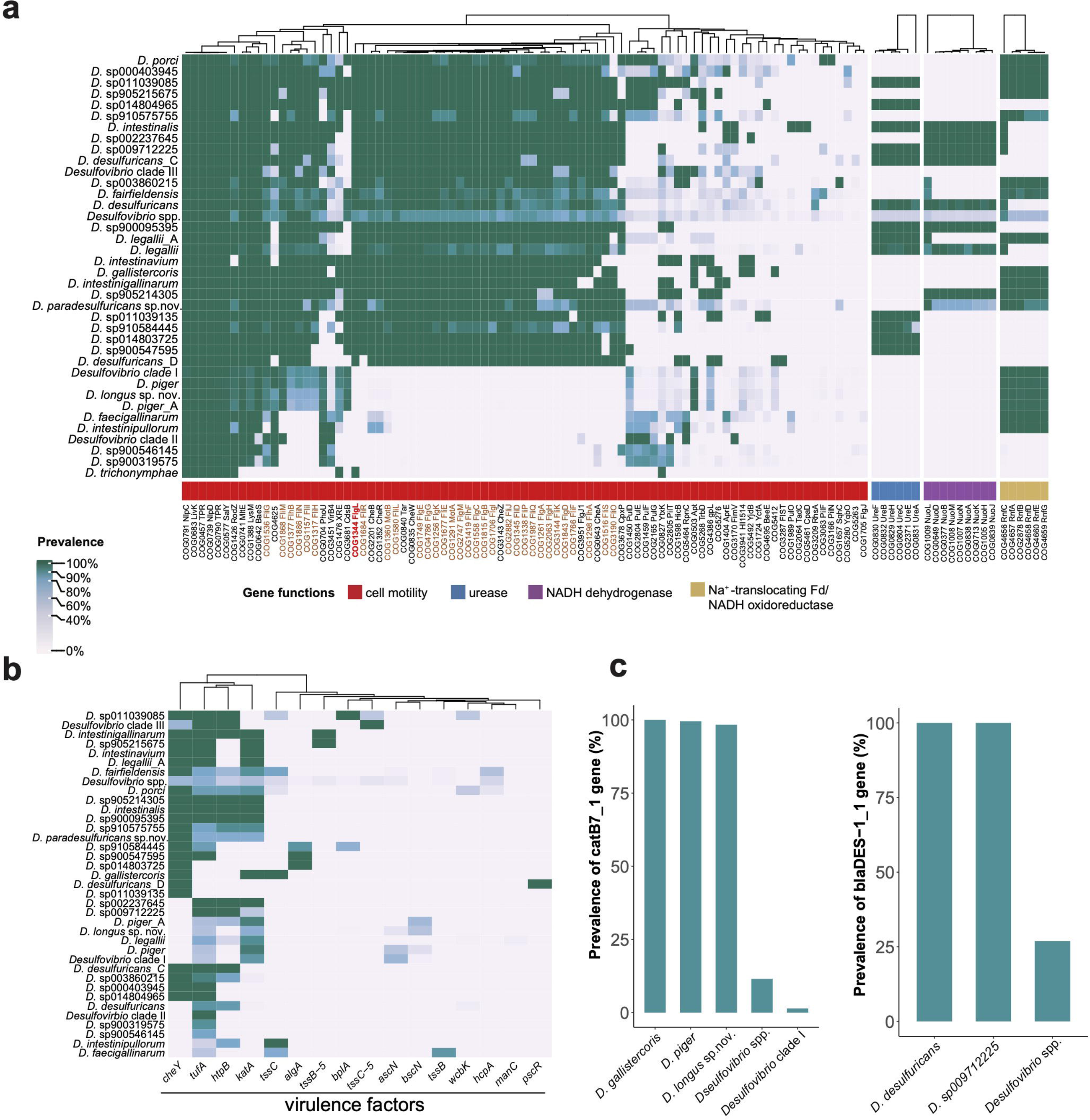
Comparison of functional potential across different *Desulfovibrio* species. **a.** The heatmap shows that the functional capabilities of *Desulfovibrio* species differ in clusters of orthologous groups (COGs) related to cell motility, urease, NADH dehydrogenase, and Na^+^-translocating Fd/NADH oxidoreductase. For cell motility, only COGs present in at least one species with a minimum prevalence of >10% were selected for the visualization. The flagellin COG is indicated in red and other flagellar-related COGs highlighted in brown. **b-c**. The heatmap and barplots show the genomic diversity in (**b**) antimicrobial resistance genes and (**c**) virulence factors. Genes were identified using ABRicate with a cutoff of >60% identify and >80% coverage. Species names follow GTDB taxonomy and represent genome-based species-level clusters. *Desulfovibrio* spp. refers to multiple *Desulfovibrio* genomes assigned to the *Desulfovibrio* genus but cannot be resolved to a known species (based on GTDB).

In addition, we identified differences in genes encoding urease, NADH dehydrogenase complex, and Na^+^-translocating Rnf complex (**Fig. 2a)**. Urease plays a role in bacterial virulence [48]. For example, urease in *Helicobacter pylori* produces ammonia at levels toxic to host epithelial cells and can neutralize stomach acidity facilitating colonization[49]. More than 95% genomes of *D. desulfuricans*, *D. legallii*, *Desulfovibrio sp910584445*, and *Desulfovibrio sp900547595* encoded the *ureABCEFH* operon required for urease synthesis and activity. In contrast, these genes were absent in *D. piger* and *D. piger_A* genomes. Furthermore, the *nuo* gene clusters (*nuoABDHJKLMN*) encoding proton-pumping type I NADH dehydrogenase was present in several species, including *D. desulfuricans*, *D. legallii*, and *D. intestinalis*, but absent in *D. porci*, *D. sp900319575* and several other species (**Fig. 2a**). This NADH dehydrogenase complex was previously identified in *Solidesulfovibrio magneticus* (formerly *Desulfovibrio magneticus*) [50]. It connects NADH oxidation to proton translocation, involving energy transfer reactions during sulfate respiration [51]. Lastly, we also detected a *rnfCDGEFB* gene cluster encoding Na^+^-translocating Rnf complex in >90% genomes of 18 *Desulfovibrio* species, including *Desulfovibrio* clade I, *D. piger*, and *D. piger_A*. The Rnf complex is homologous to the electrogenic NADH:ubiquinone reductase complex (Nqr), a sodium-pumping NADH dehydrogenase [52]. Interestingly, the *nuo* and *rnf* operon co-occurred in *Desulfovibrio sp905214305*. These results indicate that *Desulfovibrio* species encode different metabolic pathways related to energy transfer involving sulfate respiration and identify functional traits (i.e. *ure*) associated with colonization and pathogenicity.

### Antimicrobial resistance genes and virulence factors differ across *Desulfovibrio* species

To further explore disease-associated gene signals, we next investigated virulence factors and antimicrobial resistance (**Fig. 2b+c**). Virulence factors were widespread, including elongation factor *tufA*, chemotaxis protein *cheY*, catalase *katA*, and heat shock protein *htpB,* which were present in >50% of *Desulfovibrio* species (**Fig. 2b**). Meanwhile, some virulence factors were species-specific, including type VI secretion system contractile sheath small subunit *tssB*, which was prevalent in *D. faecigallinarum* (27/31) and also detected in *D. fairfieldensis* (4/117), while a type III secretion system gene (*pscR)* was detected in *D. desulfuricans*_D (1/1). Antibiotic resistome profiling also revealed several prevalent but species-specific antibiotic resistance genes (**Fig. 2c**). Chloramphenicol resistance gene *catB7* was present in *D. piger* (211/212) and highly prevalent in *Desulfovibrio longus* sp. nov. (1503/1528), while beta-lactamase resistance (*blaDES-1*) was specific to *D. desulfuricans* (42/42) and detected in *D. sp009712225* (1/1). These analyses indicate that *Desulfovibrio* species are highly diverse in their encoded virulence and antibiotic resistance genes.

To further investigate the predicted antibiotic resistance of *Desulfovibrio* strains, we performed *in vitro* antimicrobial susceptibility testing for the newly isolated *Desulfovibrio* isolates (**Supplementary Fig. 3**). Isolates from 5 *Desulfovibrio* species (*D. desulfuricans*, *Desulfovibrio paradesulfuricans* sp. nov. [i.e. *D. desulfuricnas*_A based on GTDB-tk], *D. legallii*, *D. porci* and *Desulfovibrio longus* sp. nov.) were resistant to Kanamycin (minimum inhibitory concentrations [MICs] value >64 µg/ml), while only the *D. porci* strain was resistant to Ceftazidime. Overall, antibiotic resistance of different *Desulfovibrio* isolates was highly variable, suggesting species-and strain-level variation. Not all *Desulfovibrio* strains carrying putative antibiotic resistance genes were resistant to the corresponding antibiotic, where genes may not necessarily be expressed under laboratory conditions or the gene product may not be functional. Nevertheless, the detection of putative antibiotic resistance genes can still provide useful insights to identify potential antibiotic resistance reservoirs and monitor the risk of resistance emergence.

### Distinct core and accessory genes of within-species clades associate with geography

To evaluate factors influencing within-species genomic diversity, we performed permutational analysis of variance based on accessory gene presence/absence for the major *Desulfovibrio* species (>50 available genomes). *Desulfovibrio longus* sp. nov., *Desulfovibrio piger_A*, *Desulfovibrio piger*, and *Desulfovibrio* clade I, showed highly similar core gene content with no distinct cluster based on a maximum-likelihood phylogenetic tree approach. *Desulfovibrio sp900319575* differed in core and accessory genes with distinct clusters for genomes from East Asia and the Netherlands (**Fig. 3a+b**). This was further supported by mash distance-based phylogenetic analysis (**Supplementary Fig. 4**). To investigate how geography associates with a species’ functional capacity, we identified genes enriched in Dutch and East Asian *D. sp900319575* (**Fig. 3c, Supplementary Table 7**). East Asian *D. sp900319575* enriched genes (East Asia prevalence ≥30%, Netherlands prevalence ≤10%) included multidrug resistance genes *acrB* and *mexA*, transcriptional regulatory gene *qseB* and swarming motility regulation sensor gene *rssA*. The detected enrichment of multidrug resistance genes in *D. sp900319575* within the East Asian population may be due to increased antibiotic usage [53] leading to an increase in the prevalence of resistant lineages. Meanwhile, N,N’-diacetyllegionaminic acid synthase *legI* and antitoxin *dinJ* were enriched in *D. sp900319575* within Dutch populations. Notably, *legI* is involved in legionaminic acid biosynthesis, which is incorporated into flagellin subunits [54]. Moreover, *qseB* (regulator of bacteria adherence) and *dinJ* (involved in bacterial biofilm formation) are differentially distributed within genomes from East Asia and the Netherlands, suggesting a distinct *Desulfovibrio* virulence gene repertoire. Lastly, several insertion sequence (IS) elements linked to bacterial adaptation and evolution were highly enriched in Dutch *D. sp900319575* genomes, including ISSau1, ISAsp8, and ISAzs12 (**Fig. 3c**). The most prevalent IS element was ISAsp8 (56.04%), followed by ISAzs12 (42.86%) and ISSau1 (37.36%), with much lower prevalence in East Asian genomes (ISAsp8: 3.85%, ISAzs12: 5.77%, ISSau1: 7.69%). Interestingly, these IS elements were unique to *D. sp900319575* genomes (only exception: ISSau1 was detected in one *D. sp900546145* genome). To further explore the functional context of these IS elements, we profiled and annotated their neighboring genes (within 5 kbp). Functional annotations of neighboring genes were highly diverse, including multidrug resistance and toxin-antitoxin system. These results suggest increased genome plasticity in *D. sp900319575* from East Asia population relative to Dutch populations.

**Fig. 3.**
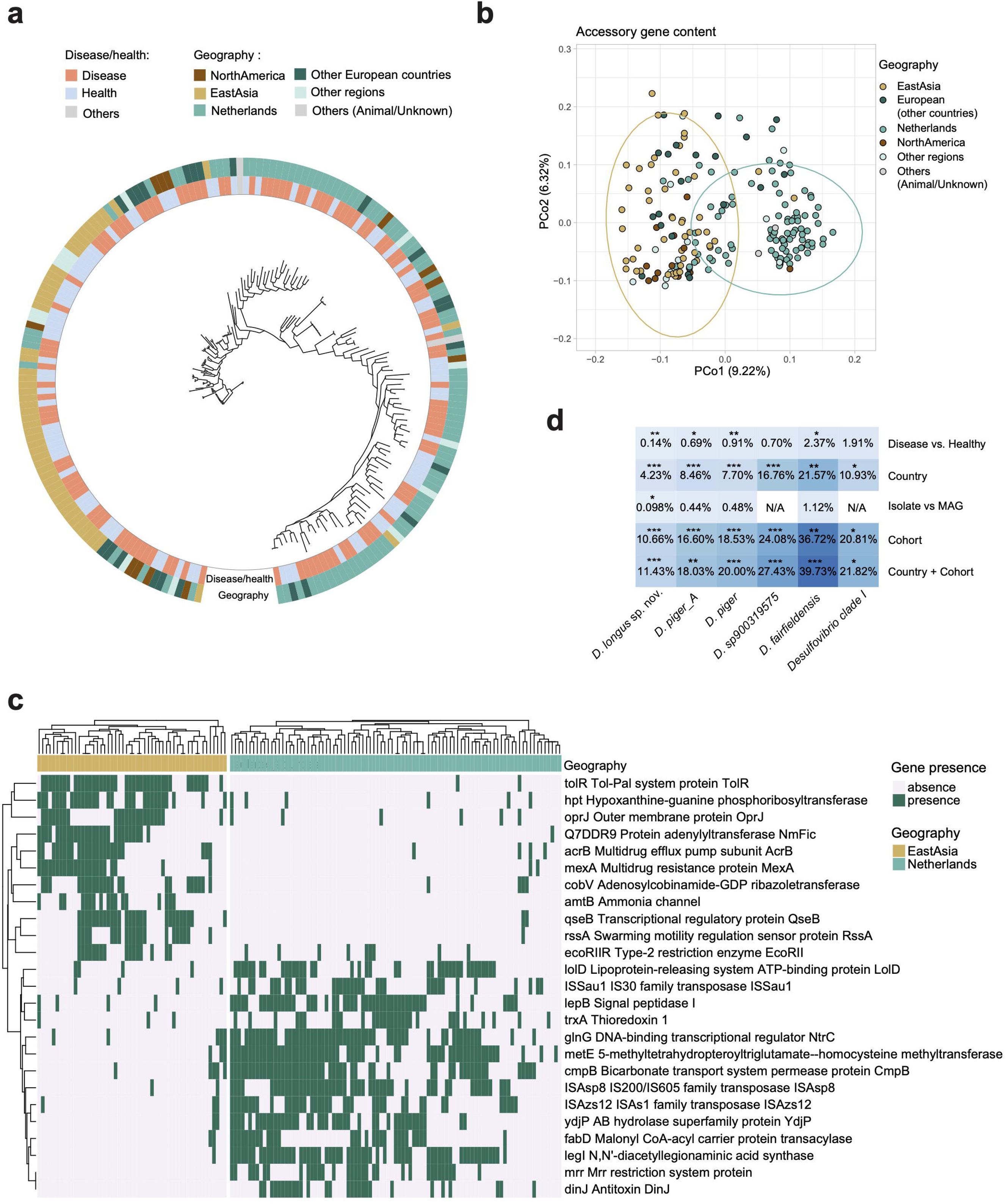
Associations of encoded gene content of *Desulfovibrio sp900319575* with geography. **a.** Maximum-likelihood phylogenetic tree based on core gene alignment of *Desulfovibrio sp900319575* genomes. Core genes were identified based on their presence in ≥ 99% of genomes of the species at ≥ 95% identity (Roary). Inner ring colors indicate host phenotypic characteristics. “Others” includes genomes from animals or unknown sources. Outer ring colors indicate geographic regions from which the genomes originate. Regions with <5 available genomes (Middle East, South America, Africa, Central Asia) were summarised as “Other regions”. **b.** PCA analysis of accessory genes within *Desulfovibrio sp900319575* genomes (Jaccard distance). Points represent *Desulfovibrio sp900319575* genomes, colours indicate host regions. **c.** Heatmap showing predicted genes (Prokka) that were enriched in the Netherlands (prevalence ≥30% in the Netherlands and prevalence ≤10% in East Asia) and East Asia (prevalence ≥30% in East Asia and prevalent ≤10% in the Netherlands), respectively. **d.** Variance in accessory gene content of major *Desulfovibrio* species explained by disease, country, genome type (isolate *vs.* MAG) and cohort (or study) of origin (PERMANOVA). Stars represent different significance level: * p ≤ 0.05; ** p ≤ 0.01; *** p ≤ 0.001. N/A indicates that genomes are only from one category of the respective variable and excluded from the analysis.

To evaluate factors associated with within-species adaptation, we performed permutational analysis of variance for the major *Desulfovibrio* species (>50 available genomes, **Fig. 3d**). Country and cohort were associated with variation in accessory gene content for *D. sp900319575* (country: R^2^=16.76%, p=0.001; cohort: R^2^=24.08%, p=0.001), *Desulfovibrio longus* sp. nov. (R^2^=4.23%, p=0.001), *D. piger_A* (R^2^=8.46%, p=0.001), *D. piger* (R^2^=7.70%, p=0.001), *D. fairfieldensis* (R^2^=21.57%, p<0.01) and *Desulfovibrio* clade I (R^2^=10.93%, p<0.05). Furthermore, disease status showed a significant but weaker association with accessory gene content in *Desulfovibrio longus* sp. nov., *D. piger_A*, *D. piger* and *D. fairfieldensis.* In addition, genome type (i.e. MAG vs isolate) was weakly associated with gene content for *Desulfovibrio longus* sp. nov. (R^2^=0.098%, p<0.05). Overall, country and cohort showed the largest effect on within-species accessory gene variation indicating species divergence.

### Flagellated CD- and CRC-associated *D. desulfuricans* and *D. legallii* activate TLR5

*Desulfovibrio* are associated with diseases, in particular IBD and CRC [1] [2]. We next identified high-level functional traits uniquely encoded by IBD- and CRC-associated *Desulfovibrio* genomes, including coenzyme and lipid transport and metabolism genes (UC, CD and CRC *vs.* healthy, Wilcoxon test, p<0.05, **Supplementary Fig. 5**). To further investigate specific disease-associated *Desulfovibrio* functional traits, we analysed IBD and CRC-associated *Desulfovibrio* COGs (**Fig. 4a+b**). In CD, 78 COGs were enriched and 15 depleted compared to healthy individuals (Fisher’s exact test; adjusted p <0.25, **Fig. 4b, Supplementary Table 8**). Toxin *hipA* (COG3550), which increases antibiotic resistance in *E. coli* [55], was significantly enriched in UC (Fisher’s exact test, adj. p=0.067, **Supplementary Fig. 6**). COGs associated with health included ribosome 30S subunit, while CD-enriched COGs were related to isoprenoid biosynthesis. Notably, urease genes *ureABCEF* were among the most significantly enriched *Desulfovibrio* operons in CD (**Fig. 4b**). Urease is a CD-associated bacterial virulence factor [49, 56] and our analyses identified a potential connection with *D. desulfuricans* and *D. legallii*. For CRC, 195 COGs showed increased prevalence in disease, including COG1111 containing putative ERCC4-related helicase MPH1, a DNA damage repair protein (**Fig. 4a, Supplementary Table 9**). The most significant gene in CRC (Fisher’s exact test; adjusted p =0.00026) was a putative MPH1-like helicase gene, which was highly prevalent in CRC and present in 509 of 2,658 *Desulfovibrio* genomes. Most (63.16%) of the identified MPH1-like genes possessed helicase core domains related to DNA repair helicases and genome stability, including PF00270 (DEAD/DEAH box helicase) and PF00271 (helicase conserved C-terminal domain). In addition, 77.78% of these genes also harbored the PF09369 domain (MrfA Zn-binding), previously reported as a DNA repair domain in *Bacillus subtilis* [57, 58]. This suggests a potential association of this gene with bacterial survival under stress. CRC-depleted genes in *Desulfovibrio* included cell shape-determining gene *mreC* (COG1792) (Fisher’s exact test, adj. p=0.00093), involved in organizing peptidoglycan synthesis and cell morphogenesis in bacteria. The depletion of *mreC* attenuates key virulence systems, i.e. SPI-1 Type III secretion system (T3SS) and flagella, in *Salmonella* [59], establishing a connection between *mreC* and bacterial pathogenicity.

**Fig. 4.**
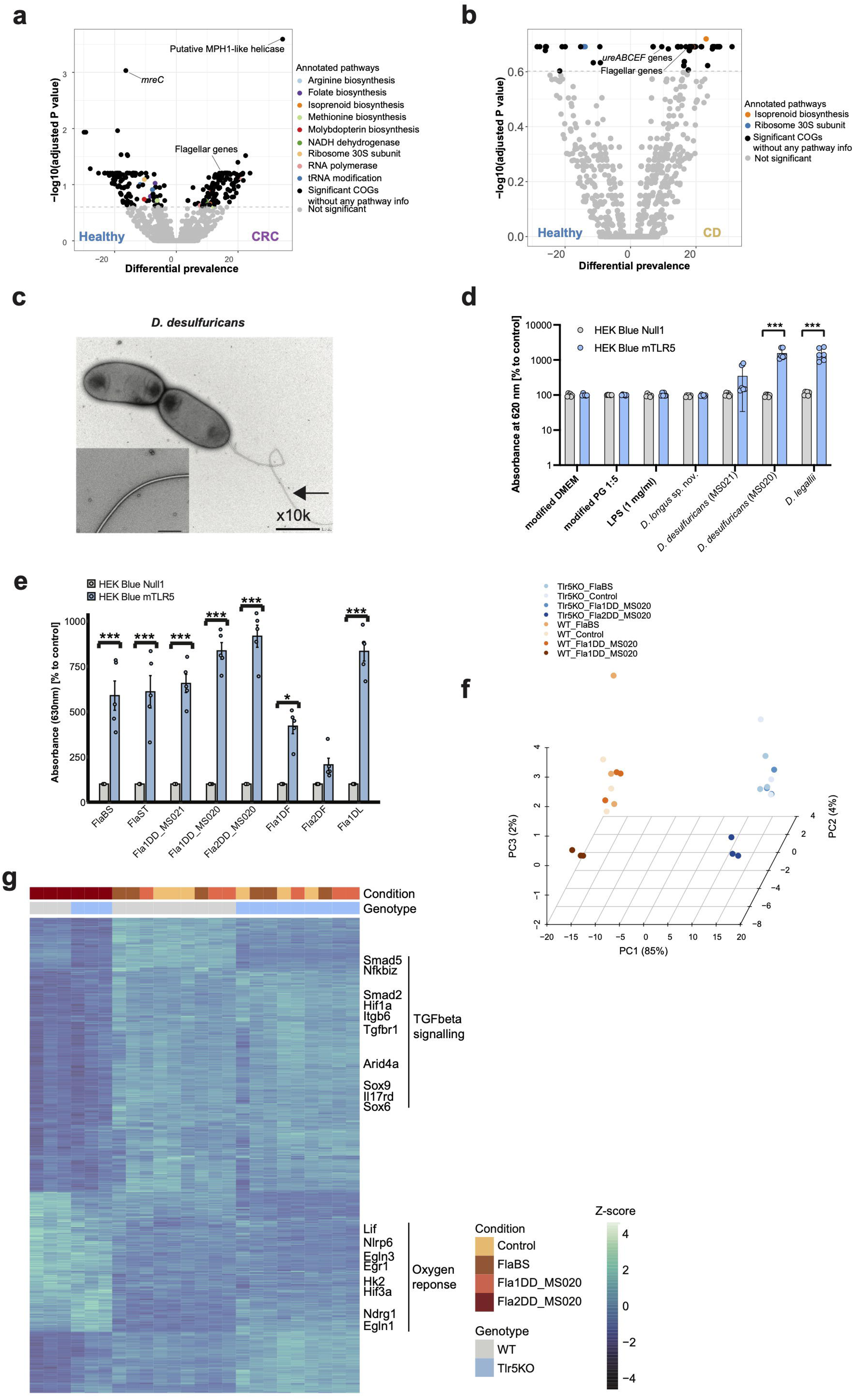
Disease-associated functional trait across *Desulfovibrio* genomes. **a** & **b**. Volcano plots showing COGs differentially enriched in CRC (**a**) and Crohn’s disease (CD) (**b**). Each point represents a COG. X-axis indicates COG prevalence in CD (or CRC) patients minus prevalence in healthy individuals. Y-axis shows log-transformed adjusted p-values (Fisher’s exact test, Benjamini-Hochberg correction for multiple testing). Dashed horizontal lines indicate significance threshold (adjusted p-value <0.25). **c.** Electron microscopy image of *D. desulfuricans* (MS020) shows visible flagella. **d.** Flagellated *D. desulfuricans* (MS020) and *D. legallii* (MS029) activate TLR5, while non-flagellated *Desulfovibrio longus* sp. nov. (MS012) showed no TLR5 activation. HEK-Blue TLR5 reporter cells were treated with bacterial culture supernatant filtrates. **e.** Recombinant flagellin proteins confirm flagellin-specific TLR5 activation. All but one of the recombinantly expressed flagellins (Fla2DD_MS020) significantly activated TLR5 (HEK-Blue TLR5 reporter cell line, linear mixed-effects model with Sidak comparison, stars represent significance level: * p<0.05; **** p<0.0001. The concentrations for FlaBS and FlaST are 0.5ng/ml and the concentrations for all recombinant *Desulfovirbio* flagellins are 1000 ng/ml. Abbreviations of the flagellins on the x-axis are defined as follows: FlaBS, flagellin from *Bacillus subtilis;* FlaST: flagellin from *Salmonella typhimurium*; Fla1DD_MS021, flagellin 1 from *D. desulfuricans* (isolate MS021); Fla1DD_MS020, flagellin 1 from *D. desulfuricans* (isolate MS020); Fla2DD_MS020, flagellin 2 from *D. desulfuricans* (isolate MS020); Fla1DF: flagellin 1 from *D. fairfieldensis* (isolate MS026); Fla2DF: flagellin 2 from *D. fairfieldensis* (isolate MS026); Fla1DL: flagellin 1 from *D. legallii* (isolate MS029). **f**. Fla2DD_MS020 flagellin induces distinct gene expression patterns in murine small intestinal organoids. PCA analysis based on bulk RNAseq of flagellin-treated organoid experiments. WT = wild type organoids, Tlr5KO = TLR5 knock-out organoids. Unstimulated organoids were used as negative controls. **g.** Differentially expressed genes induced by Fla2DD_MS020 include upregulation of TGF-beta signaling and downregulation of genes involved in oxygen response (Wald test implemented in R package DESeq2, adjusted p <0.05).

Overall, 62 disease-enriched COGs were shared between CD and CRC, with 22 of these being flagellar-related, including flagellar basal body rod gene *flgC*, flagellar basal body-associated gene *fliL* and flagellin gene *flgL* (**Fig. 4a+b**). Flagellins have been linked to inflammation via their ability to stimulate innate immune receptor TLR5 [60, 61]. To investigate immune system activation in connection with flagellated *Desulfovibrio* spp., we profiled flagellar-related genes in our clinical isolates. We identified disease-enriched flagellar-related genes in all *D. desulfuricans*, *D. fairfieldensis* and *D. legallii* isolates, while these were absent in *D. piger* isolates. Transmission electron microscopy (TEM) images confirmed visible flagella in *D.desulfuricans* (MS020)*, D. fairfieldensis* (MS026) and *D. legallii* (MS029), while no flagellation was observed in *Desulfovibrio longus* sp. nov. (MS012), as expected (**Fig. 4c, Supplementary Fig. 7**). These results are consistent with a previously observed flagellar structure in a *D. legallii* strain [28]. N- and C-terminal functional domains responsible for TLR5 activation were present in flagellin genes *flaB2*, *flaB3* and flagellar filament 33 kDa core gene in all *D. desulfuricans* and *D. fairfieldensis* and most *D. legallii* genomes (28/30), suggesting a central role of *flaB* subunits. Flagellin gene structure predictions support that the *flaB* subunits of *D. desulfuricans* isolates harbor orthologs of D0 and D1 domains of *Salmonella*-derived flagellin (FliC), a known TLR5 activator [60] (**Supplementary Fig. 8**). We validated TLR5 activation by *D. desulfuricans* and *D. legallii* (**Fig. 4d**). For this, HEK-Blue TLR5 reporter cells were stimulated with bacterial culture supernatant filtrates, experimentally confirming TLR5 activation by flagellated *Desulfovibrio*. Overall, these results link flagellated *Desulfovibrio* to immune system activation in CD and CRC.

### *D. desulfuricans* flagellin treated organoids suppress TGF-beta signaling

To further investigate the immune response induced by these flagellins, we recombinantly expressed and purified *flaB2*, *flaB3* and flagellar filament 33 kDa core protein genes from *D. desulfuricans* (Fla1DD_MS020, Fla2DD_MS020, Fla2DD_MS021), *D. legallii* (Fla1DL) and *D. fairfieldensis* (Fla1DF and Fla2DF). All but one flagellin (Fla2DF) significantly activated TLR5 (HEK Blue reporter cell lines, **Fig. 4e**). We further profiled immune system interactions involving *Desulfovibrio* flagellins using murine small intestinal organoids (*Tlr5* knocked-out vs wild type organoids). Image-based analysis of all treated organoids confirmed no significant differences in growth between control flagellins (FlaBS and FlaST from *Bacillus subtilis* and *Salmonella typhimurium*, respectively) and *Desulfovibrio*-derived proteins (**Supplementary Fig. 9**). This indicates that these proteins do not induce acute toxic effects on epithelial cells and transcriptional differences were not due to skewed differentiation or growth of the organoids.

We subsequently exposed wild-type and TLR5 knockout mouse organoids to the two *Desulfovibrio* flagellins from the same isolate (Fla1DD_MS020 and Fla2DD_MS020), including FlaBS as a positive control, and performed bulk RNA-seq. Fla2DD_MS020–treated organoids exhibited distinct gene expression profiles compared to other flagellins (**Fig. 4f**). As expected, FlaBS induced an inflammatory response characterised by increased expression of TNF, NF-κB, and antimicrobial peptide genes (**Supplementary Fig. 10)** [62, 63]. While both Fla1DD_MS020 and Fla2DD_MS020 lacked the same canonical TLR5 signature, Fla1DD_MS020 stimulation resulted in reduced expression of several genes via TLR5 signaling, including *Trib3* (a regulator of stress and inflammatory signaling), and multiple solute carrier genes involved in cellular transport. Interestingly, Fla2DD_MS020 induced a robust TLR5-independent transcriptional signature with a significant downregulation of the TGF-beta signaling pathway, including *Tgfbr1*, *Smad2/5*, and *Sox6/9* genes, while oxygen response pathways were upregulated, involving *Nlrp6*, *Egr1* and *Egln3* expression (**Fig. 4g**). No compensatory activation of other TLRs was detected **(Supplementary Fig. 11)**. While flagellins can also activate intracellular receptors of the nucleotide binding oligomerization domain [NOD]-like receptors (NLR) family, only *Nlrp6* showed strong upregulation by Fla2DD_MS020 (Wald test; wild type organoids: Fla2DD_MS020 *vs.* control, adjusted p = 5.17674e-43; TLR5 knock-out organoids: Fla2DD_MS020 *vs.* control, adjusted p = 6.69439e-47. **Supplementary Fig. 12)**. NLRP6 is an intracellular inflammasome activator that is upregulated in response to TNF as well as microbial and viral triggers [64], however, no expression changes of IL-1β, IL-18, and ASC/Pycard were observed, suggesting that increased *Nlrp6* transcription did not lead to inflammasome activation. Overall, these results demonstrate that flagellated *Desulfovibrio* are active modulators of the innate immune response with distinct immune signatures compared to canonical flagellins from *B. subtilis* and *S. typhimurium*.

### Dissimilatory sulfate reduction is highly conserved across *Desulfovibrio* spp

Sulfur metabolism is one of the key pathways in *Desulfovibrio* that has been hypothesised to play a role in disease. We next compared sulfur metabolism capabilities across *Desulfovibrio* genomes. For this, we curated a database of sulfur metabolism genes based on the Kyoto Encyclopedia of Genes and Genomes (KEGG) pathways [65] and previous studies [16, 25, 66, 67]. There are five major pathways involved in H_2_S production: dissimilatory and assimilatory sulfate reduction, cysteine degradation, tetrathionate metabolism and taurine metabolism (**Supplementary Table 10**). The majority of *Desulfovibrio* genomes (81.34%) encoded the complete dissimilatory sulfate reduction pathway (hmmscan, gene hits with e-value ≤ 1E-10). Further, >99% of the genomes contained ≥80% of the pathway genes, where an incomplete pathway was associated with lower genome completeness (**Supplementary Fig. S13**). Overall, this indicates that dissimilatory sulfate reduction is highly conserved across *Desulfovibrio* species.

Furthermore, we identified several taurine metabolism gene clusters. One of these clusters was associated with taurine metabolism to isethionate, which is subsequently converted to sulfite (**Supplementary Fig. 14**), which was previously reported in *D. desulfuricans* DSM 642 and *Oleidesulfovibrio alaskensis* G20 (formerly *Desulfovibrio alaskensis* G20) [67]. While many taurine metabolism genes (e.g. taurine pyruvate aminotransferase [*tpa*] and sulfoacetaldehyde acetyltrasferase [*xsc*]) were well conserved across *Desulfovibrio* spp. (**Supplementary Fig. 15**), we identified species-specific differences in genes encoding alanine dehydrogenase (*ald)*. All *D. desulfuricans*, *D. legallii_A*, *D. intestinalis*, and *D. sp011039135* genomes encoded *ald*, while the gene was absent in *D. piger*, *D. piger*_A, *Desulfovibrio* clade I, *Desulfovibrio* clade II and clade III. These results suggest that while *Desulfovibrio* spp. shared core taurine metabolism pathway, variation in *ald* suggests species-specific difference in processing taurine-derived intermediates.

While dissimilatory sulfate reduction gene prevalence was comparable between individuals with IBD or CRC and healthy individuals (Fisher’s exact test, adjusted p>0.25), other sulfur metabolism genes were enriched in disease, in particular *ald* gene for supporting taurine metabolism and *ttrBCA* genes responsible for tetrathionate respiration (enriched in CRC and CD, Fisher’s exact test, adjusted p<0.25, **Supplementary Fig. 16**). Conversely, *yhaO*, involved in cysteine degradation, showed reduced prevalence in CD and CRC compared to healthy individuals. Overall, this suggests that other H_2_S pathways are encoded by IBD- and CRC-associated *Desulfovibrio* spp..

### Investigating H_2_S production capacity of the IBD microbiome

Next, we investigated the overall IBD microbiome capacity for H_2_S production. We searched for sulfur metabolism genes in all MAGs in a deeply sequenced IBD cohort [68] (hmmscan, gene hits with e-value≤1E-10). Dissimilatory sulfate reduction was encoded by 40 of 54 (74.07%) *Desulfovibrio* MAGs spanning 7 known and one uncharacterized species (**Supplementary Table 11**). In addition, 6 *Mailhella* (37.5%) MAGs encoded 80% of the pathway genes, where all *Mailhella* genomes lacked sulfate adenylyltransferase (sat/met3). The tetrathionate pathway was more widely distributed and identified in 41 MAGs from 9 genera: *Desulfovibrio*, *Gordonibacter*, *Citrobacter*, *Proteus*, *Eggerthella*, *Escherichia*, *Rubneribacter*, *Morganella* and *Turicimonas* (**Fig. 5a+b**). Overall, 443 MAGs from 16 genera, including *Escherichia* and *Citrobacter*, encoded seven genes involved in assimilatory sulfate reduction (*cysC*, *cysD*, *cysH*, *cysI*, *cysJ*, *cysN*, *cysNC*) and only lacked sulfate adenylyltransferase (*sat*) and sulfite reductase (ferredoxin) (*sir*) (**Fig. 5a+b**). As *cysD* and *cysN* together perform the same function as *sat*[69, 70], and *cysIJ* are alternatives for *sir*[71, 72], these bacteria still maintain the genomic capacity required for sulfide production through the assimilatory sulfate reduction pathway. While sulfide is typically directly incorporated into cysteine biosynthesis [73, 74] and does not result in free H_2_S [75], cysteine can be further degraded, leading to an indirect contribution of free H_2_S. Overall, our genomic analysis revealed several bacteria that encode the complete tetrathionate metabolism, assimilatory and dissimilatory sulfate reduction pathway, identifying these pathways as potential H_2_S sources (including relevant precursors) in this IBD cohort.

**Fig. 5.**
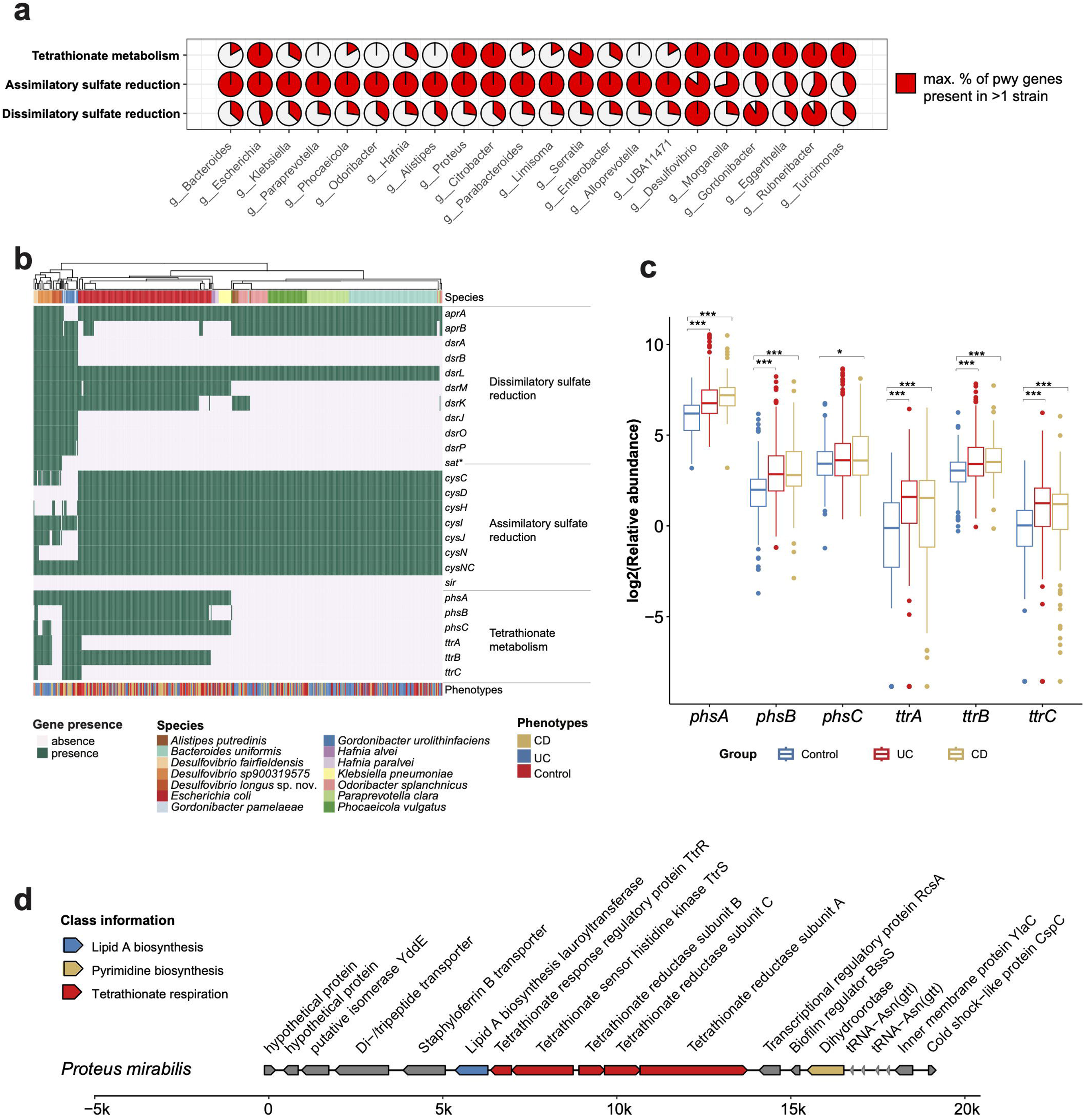
H_2_S pathway-encoding bacteria in IBD. **a.** Three pathways involved in H_2_S production (dissimilatory sulfate reduction, assimilatory sulfate reduction and tetrathionate metabolism) were encoded by different genera (FedericiS_2022 cohort [68]). Only genera, where 100% of the necessary pathway genes were present in at least one genome, are shown. Assimilatory sulfate reduction: *cysD* and *cysN* are functionally equivalent, *sat* and *cysIJ* are alternatives for *sir*, therefore strains lacking *sat* or *sir* but encoding *cysD*, *cysN*, and *cysIJ* were still classified as encoding the complete assimilatory sulfate reduction pathway. **b**. The heatmap shows the presence of genes related to three pathways involved in H_2_S production (dissimilatory sulfate reduction, assimilatory sulfate reduction and tetrathionate metabolism) in species with >3 MAGs encoding at least one complete pathway. \**sat* is involved in dissimilatory and assimilatory sulfate reduction according to KEGG modules M00596 and M00176. **c.** Genes involved in tetrathionate metabolism were identified at community level with significantly increased abundance in UC and CD patients compared to controls. Wilcoxon test: * p < 0.05; ** p<0.01; *** p<0.001. **d.** Genomic context of the *ttrBCA* operon in the *Proteus mirabilis* MAGs. COG pathway information is indicated by different colors.

Due to the lack of a universally conserved gene sets and the presence of many alternative enzymes in bacterial taurine metabolism and cysteine degradation, we did not assess completeness of these pathways. Instead, we explored these pathways at the microbial community level to identify their associations with IBD. Several taurine metabolism genes were significantly increased in IBD patients compared to healthy individuals (adj. p<0.05, **Supplementary Fig. 17**), including enzymes that catalyse the metabolism of taurine to isethionate with subsequent conversion to sulfite (*tpa*, *sarD*, *islA* and *islB*). In contrast, phosphotransacetylase (*pta)*, which converts acetyl-phosphate to acetate, a short-chain fatty acid important for human epithelial integrity [76], showed reduced abundance in IBD (adj. p<0.05, **Supplementary Fig. 17**). Moreover, taurine dehydrogenase large subunit *tauY* abundances were increased in UC but decreased in CD compared to healthy individuals. These results indicated an association of changes in taurine metabolism gene abundances in IBD connected to acetate production. Similarly, most genes related to cysteine degradation are more abundant in IBD patients compared to healthy individuals (adj. p<0.05, **Supplementary Fig. 18**). In particular, the major cysteine-degrading genes (*dcyD*, *yhaM* and *sseA*) were increased in IBD samples. Moreover, *sseA* has been found to be associated with bacterial virulence [77]. These results indicate an increase in bacteria encoding H_2_S-associated taurine and cysteine degradation pathways in IBD patients.

The majority of MAGs (86.49%) encoding tetrathionate metabolism were recovered from IBD patients. Interestingly, these MAGs encoded the *ttr* genes, responsible for tetrathionate respiration, which has been shown to confer growth advantages in pathogenic bacteria during inflammation [78]. Indeed, the abundance of all genes involved in this pathway was significantly higher in IBD compared to healthy individuals (exception: *phsC* in UC *vs.* healthy, p=0.074, Wilcoxon, all other adj. p<0.05, **Fig. 5c**). To further investigate gene activity related to tetrathionate metabolism, we analysed IBD metatranscriptomic data [79]. In CD patients, all but one tetrathionate metabolism gene showed increased expression compared to non-IBD controls (exception: *phsB* in CD *vs.* non-IBD, p=0.46, Wilcoxon, all other adj. p<0.05, **Supplementary Fig. 19**). For UC patients, *phsA*, *phsC* and *ttrB* were expressed in most of the samples, while only *ttrA* showed a significant increase in UC *vs.* non-IBD, which may be due to smaller UC sample numbers. MAGs encoding the complete tetrathionate metabolism pathway included 5 *Proteus* MAGs from 3 species (*P. mirabilis*, *P. vulgaris*, *P. sp003144505*). *Proteus mirabilis,* a known CD-associated pathogen [80], contained *ttrRS* and *ttrBCA* gene clusters (**Fig. 5d**), which can confer competitive growth advantages [80]. Overall, these results indicate that tetrathionate metabolism abundance is increased and transcriptionally active in CD and identifies other potential new players encoding disease-associated H_2_S production pathways.

## Discussion

While the number of culturable bacteria has tremendously increased over the past years, the isolation of *Desulfovibrio* strains remains challenging, greatly limiting our ability to study their role in human health. To close this gap we combined large-scale bioinformatic profiling across metagenomic human samples with targeted isolation efforts to establish a comprehensive *Desulfovibrio* database in connection with a collection of representative isolates to identify disease-associated functional traits.

We identified disease-associated flagellar genes in *D. desulfuricans*, *D. fairfieldensis* and *D. legallii* and confirmed TLR5 and strain-specific immune system activation, providing a potential link to IBD and CRC pathogenesis. In particular, we observed that flagellin from *D. desulfuricans* strongly suppressed TGF-beta signaling in murine small intestinal organoids. TGF-beta is a pleiotropic transcription factor with key roles in maintaining mucosal tolerance by promoting anti-inflammatory and tissue repair programs, specifically at the epithelial layer, where TGF-beta signaling is required for epithelial cell regeneration in response to damage [81]. Furthermore, TGF-beta promotes adaptive immune tolerance through persistence of intraepithelial T cells in combination with retinoic acid (RA) [82] and is essential for establishing intraepithelial CD8αα [83] and CD4 T cells [84]. The intestinal innate immune compartment also requires TGF-beta to regulate monocyte/macrophage recruitment and tissue-specific imprinting [85]. In human intestinal epithelial cells, TGF-beta and RA signaling promote tolerogenic CD103+ DCs [86] and suppressed TGF-beta signaling in CD patients is associated with chronic inflammation [87], while TGF-beta1 deficiency causes severe early onset IBD [88, 89]. Thus, the observed downregulation of TGF-beta signaling induced by *D. desulfuricans* flagellin (Fla2DD_MS020) may lead to disruptions of both innate and adaptive immune tolerance. Interestingly, the observed effects were independent of TLR5. Thus, we hypothesize that this protein binds to a different surface receptor, in addition to TLR5, which remains to be identified.

Furthermore, the virulence factor urease was encoded by *D. desulfuricans* and *D. legallii* and highly enriched in CD patients. Urease is implicated in CD development through nitrogen compound circulation [56], suggesting a potential connection between *Desulfovibrio* nitrogen metabolism through urease activity in the gut and CD development. Importantly, expression of *Desulfovibrio* nitrogenase genes (urea uptake system) in patient stool samples suggests that nitrogen fixation is active in the human gut [28]. Together, these results indicate species-specific pathogenic associations of *Desulfovibrio* with inflammatory diseases.

Current hypotheses suggest a connection between *Desulfovibrio* and disease through H_2_S production. While *Desulfovibrio* are mainly known for dissimilatory sulfate reduction, *Desulfovibrio*-associated genes of this pathway showed no differences in disease, however, genes involved in tetrathionate-related H_2_S production were highly abundant in the IBD gut microbiome compared to healthy individuals. Importantly, we identified additional disease-associated bacteria with the genomic potential to produce H_2_S via tetrathionate metabolism, including *Proteus mirabilis* and *Morganella morganii*. Both species have been implicated in IBD and CRC, where *P. mirabilis* induces pro-inflammatory cytokines and activates inflammation-related pathways in CD, including NOD-like receptor signaling [80], and *M. morganii* is enriched in IBD and CRC, producing genotoxic indolimines and exacerbating colon tumorigenesis in gnotobiotic mice [90]. Interestingly, we exclusively detected *M. morganii* MAGs in IBD patients. Overall, our results motivate future studies of potential disease mechanisms through H_2_S for these bacteria, including transcriptomic analyses to delineate bacterial H_2_S contribution in disease.

Overall, we were able to generate important insights into *Desulfovibrio* genomic diversity and identified disease-associated functional traits. However, some limitations apply as well. Firstly, we captured a substantial amount of *Desulfovibrio* spp. genetic diversity across the world, investigating samples and strains from 174 microbiome studies spanning 32 countries. However, as many countries are still understudied and lack detailed microbiome cohort studies, we expect that additional *Desulfovibrio* species will emerge as more cohort studies become available. Furthermore, most gut microbiome studies rely on stool due to ease of sample collection and high bacterial content. However, qPCR measurements have previously indicated high prevalence of *D. desulfuricans* and *D. vulgaris* in endoscopic rectal biopsies [6, 91]. Stool and other intestinal regions may differ in community composition as bacteria may adapt to the respective physiological environment [92, 93]. Secondly, we reported putative H S-producing bacteria by identifying their genomic repertoire and investigating KEGG and previously curated pathways. This does not account for alternative, functionally equivalent enzymes that may be present in some bacteria and synergistic bacterial species. Further, H_2_S measurements are currently lacking for most IBD cohorts and accurate H_2_S quantification remains challenging due to its high volatility and instability, particularly for fecal samples. Further method development is required to disentangle H_2_S production by different microbial and host-related pathways. Lastly, while we included preliminary characterization of the isolates from three putative *Desulfovibrio* species, a comprehensive taxonomic description is required for formal validation of the proposed names and will be addressed in future work.

## Acknowledgements

We would like to thank Tom Clavel (RWTH University Hospital) and Suzanne Devkota (Cedars-Sinai Medical Center) for making *Desulfovibrio* strains available. Moreover, we would like to thank Klaus-Peter Janssen (TUM Klinikum Rechts der Isar) and Tobias Schwerd (LMU Pediatric Gastroenterology and Hepatology) for sharing stool samples for our *Desulfovibrio* isolation efforts. We further want to acknowledge support from the Deutsche Forschungsgemeinschaft (DFG, German Research Foundation) – project number 395357507 (SFB 1371, Microbiome Signatures) to MS and DH. AJ, KC and MY are supported by BMBF 16LW0432, DFG TRR353-471011418, DFG/CRC1371-395357507 and the DFG-504986538 grants. R. G is supported by NWO Vidi grant VI.Vidi. 233.079.

## Materials and Methods

### Human *Desulfovibrio* isolates

Detailed information for all isolates can be found in **Supplementary Table 5**.

Ten strains were purchased from public culture collections, including the German Collection of Microorganisms and Cell Cultures (DSMZ), Culture Collection University of Gothenburg (CCUG) and Collection de Souches de l’Unité des Rickettsies (CSUR). In addition, two *Desulfovibrio* isolates were kindly shared by other labs (Suzanne Devkota, Cedars-Sinai Medical Center [94], and Tom Clavel, University RWTH University Hospital [95].

Furthermore, we used stool samples from a pediatric CD study (EEN study [96]) approved by the Ethics committee of the Ludwig-Maximilians University of Munich (approval No. 17-801; German Clinical Trials Register Accession No. DRKS00013306, date registration 19.03.2018) and a CRC cohort described in [97] (Klaus Peter Janssen; Ethikvotum (375/16S)) approved by the Ethics committee of the School of Medicine and Health, TUM (2022-275-S-DFG-KK) to obtain further isolates. For this, 1 g of frozen fecal material (20 % glycerol) was transferred into 10 ml of anaerobe modified liquid Postgate medium 2 (0.5 g K_2_HPO_4_, 1.0 g NH4Cl, 1.0 g Na2SO4, 0.1 g CaCl2 x 2 H2O, 2.0 g MgSO4 x 7 H2O, 2.0 g Na-DL-lactate, 1.0 g Yeast extract, 0.5 ml Na-resazurin (0.1 % w/v), 0.5 g FeSo4 x 7 H2O, 0.1 g Na-thioglycolate, 0.1 g Ascorbic acid) in Hungate tubes previously sparged with N_2_. From this, a series of dilutions (1:10) in modified liquid Postgate medium 2 was created and 10 µl of all steps was used to streak bacteria on modified Postgate medium 2 agar (15 g/l agar), Columbia blood agar plates (7% sheep blood) and Devkota agar [94] (10 g Pancreatic digest of Casein, 10 g Peptic digest of animal tissue, 1 g Dextrose, 2 g Yeast extract, 5 g Sodium Chloride, 0.1 g Sodium bisulfite, 5 g Pyruvate, 5 g Taurine, 0.5 g Ferric ammonium citrate, 14 g Agar, 1 ml Hemin stock (0.5 g/ 10 ml NaOH 1N), 0.5 ml Vitamin K stock (1g/ 100 ml absolute ethanol)). Agar plates and tubes were incubated under anaerobic conditions (5 % CO2, 5 % H2, 90 % N2, 37 °C) and monitored for growth and the appearance of black discoloration, indicating formation of iron sulfide from hydrogen sulfide produced by the desired bacteria with iron in the growth media. Blackened liquid samples or colonies were processed further by passaging, switching between liquid medium and solid agar plates. Passages of interest were verified for the presence of *Desulfovibrio* by PCR, using 1 µl of bacterial solution in 19 µl of Mastermix (per sample 10 µl PCR-grade ddH2O, 10 µl DreamTaq Green PCR Master Mix 2x, 0.1 µl of each 100 µM primer CCGTAGATATCTGGAGGAACATCAG and ACATCTAGCATCCATCGTTTACAGC from Sigma Aldrich), with initial denaturation at 95 °C for 3 min, 30 cycles of denaturation (95 °C, 15 s), primer annealing (58 °C, 15 s) and extension (72 °C, 15 s) and a final extension at 72 °C for 1 min. Samples were separated by gel electrophoresis on a 2 % agar gel in 1X TAE buffer at 100 V and checked for bands at 135 bp. PCR positive colonies showing uniform growth on agar were further verified via 16S rRNA PCR analysis, using the same settings as before with primers AGAGTTTGATCCTGGCTCAG and GGTTACCTTGTTACGACTT from Sigma Aldrich, with initial denaturation at 95 °C for 3 min, 30 cycles of denaturation (95 °C, 20 s), primer annealing (55 °C, 30 s) and extension (72 °C, 90 s) and a final extension at 72 °C for 3 min. Samples were separated by gel electrophoresis on a 1 % agar gel in 1X TAE buffer at 100 V and processed according to manufacturer’s recommendations of the Mix2Seq Kit by Eurofins for sequencing. Results were identified by sequence alignment against public databases (EzBioCloud and NCBI BLAST). Samples positive for *Desulfovibrio* were restreaked on blood agar three times, continually choosing single colonies to guarantee the pureness of bacteria. Finally, bacteria were grown in liquid medium and frozen in either 5% DMSO or 20% glycerol. After extracting whole-genomic DNA from individual colonies, paired-end 150-bp read sequencing was performed using an Illumina NovaSeq.

Further *Desulfovibrio* isolates were obtained from stool samples from the Lifelines Dutch microbiome cohort and the 1000IBD cohort (University Medical Center Groningen Medical Ethical Committee: IRB numbers 2017.152, 2016/424 [GE-ID], 2014/291 [Rise-Up]) and two healthy US donors from OpenBiome (ethics approval MIT - COUHES protocol #1612797956) [98]. Broad culturing was performed with a rich medium, (Brain Heart Infusion, Bacto 237300), followed by whole-genome sequencing.

### DNA extraction for Oxford nanopore sequencing, library preparation and sequencing

DNA extraction mainly followed the protocol from Harju et al. [99]. Briefly, frozen cell pellets were suspended in TE buffer. After centrifugation, the sample was resuspended in 400 μl BnG lysis buffer, placed in a dry ice bath for 2 minutes, transferred to a 99°C heat block for 2 min, and vortexed for 30 seconds. Genomic DNA was further extracted with Phenol-Chloroform-Isoamyl alcohol mixture (25:24:1 [v/v/v]), phenol, and Chloroform. Afterwards, 15 μl RNase A (10 mg/ml) and 30 μl proteinase K (20 mg/ml) were added to remove contamination. Then DNA was precipitated by adding 100 μl 3 M sodium acetate (pH 5.2) and 900 μl 96% ice-cold ethanol ([v/v], pure). Quality of isolated genomic DNA was checked by gel electrophoresis, and concentration was quantified using a nanodrop (Implen NanoPotometer).

DNA libraries were prepared for long-read sequencing with the ONT Rapid Sequencing Kit SQK-RBK004 according to the manufacturer’s protocol and sequenced with the ONT MinION sequencer using R 9.4 flow cells. The sequencer was controlled with the MinKNOW v21.05.12 software. Sequencing runs were scheduled for 72 hours. After sequencing, basecalling was performed using guppy v5.0.12 (ONT) to obtain fastq files.

### *Desulfovibrio* isolate assembly

For Illumina sequencing, high-quality reads were obtained by trimming adapters and removing low-quality reads using Trim Galore [100] with parameters “--phred33 –quality 30 –stringency 5 –length 10”. For ONT sequencing, adapter removal was performed using Porechop (https://github.com/rrwick/Porechop/), followed by filtering short reads (<2kb) and removing low-quality reads (quality score <10) using NanoFilt [101]. Unicycler was used for genome assembly [102]. Hybrid assembly was done using parameters “-1”, “-2” and “-l” for the isolates with ONT and Illumina reads. For isolates with only Illumina data, parameters “-1” and “-2” were used to specify paired-end short reads.

### Retrieval of *Desulfovibrio* genomes from public databases

*Desulfovibrio* MAGs and isolate assemblies were retrieved from the following public repositories: (1) NCBI RefSeq (https://www.ncbi.nlm.nih.gov/refseq/); (2) Unified Human Gastrointestinal Genome (UHGG) [42]; (3) the Human Reference Gut Microbiome (HRGM) [103]; (4) IMG/M [104]; and (5) MGnify (http://www.ebi.ac.uk/metagenomics) [105]. The respective metadata information was also downloaded from the databases.

### Retrieval of *Desulfovibrio* genomes from MetaRefSGB database

For MetaRefSGB and the associated MetaPhlAn markers database (version Jun23_CHOCOPhlAnSGB_202403), MAGs were recovered from >100k metagenomic samples and grouped into species-level genome bins (SGBs, further details in [40]). Each SGB comprises genomes that share at least 95% ANI. Genomes with completeness >90% and <5% contamination that were annotated as *Desulfovibrio* were retrieved for this study. Taxonomic labels were reassigned using GTDB-tk (version 2.1.1) [41] and only genomes that were classified as *Desulfovibrio* spp. were retained. Genome-associated metadata was obtained from the curatedMetagenomicData resource [106], whenever possible, and otherwise sourced directly from the original publications or datasets in which the genomes were first reported.

### Metagenomic data processing, assembly and MAG identification

All public datasets we used for *Desulfovibrio* MAG construction and identification are listed in **Supplementary Table 2**. Downloaded metagenomic raw reads were processed by removing adapters and trimming sequences using Trim Galore [100] with the following parameters: --phred33 –quality 30 –stringency 5 –length 10. Then Kneaddata (https://github.com/biobakery/kneaddata) was used for quality checking, quality filtering and host sequences decontamination, by setting “--trimmomatic-options HEADCROP:15 SLIDINGWINDOW:4:15 MINLEN:50” and default parameters. High-quality reads were assembled using Megahit [107] with default parameters for each metagenomic sample. MAGs were identified based on three binning algorithms: Maxbin 2 [108], MetaBAT 2 [109] and CONCOCT [110]. Contig length with ≥1,000 bp were used for Maxbin2 and CONCOCT binning. For MetaBAT 2 binning, the minimum contig length was set as 1,500 bp. MAGs produced from the three binning methods were further refined using metaWRAP ‘Bin_refinement’ module [111].

### Identification of *Desulfovibrio* genomes in the Dutch microbiome cohort

The cohort is described in detail in [39]. Briefly, the Dutch Microbiome Project (DAG3) cohort includes 8,208 individuals from the Northern Netherlands (age-range 8 84 years, 57.4% female, 99.5% Dutch European ancestry). Fecal samples were collected between 2015-2016 and microbial DNA was isolated (QIAamp Fast DNA Stool Mini Kit, Qiagen - Germany). Library preparation for samples with total DNA yield <200 ng (as determined by Qubit 4 Fluorometer) was performed using NEBNext® Ultra™ DNA Library Prep Kit for Illumina, while libraries for other samples were prepared using NEBNext® Ultra™ II DNA Library Prep Kit for Illumina®. Metagenomic sequencing was performed (Illumina HiSeq 2000 platform) to generate approximately 8 Gb of 150 bp paired-end reads per sample (mean=7.9Gb, sd=1.2Gb). Preprocessing of metagenomic raw reads included adapter trimming using BBduk from the BBMap package v.38.93 (https://sourceforge.net/projects/bbmap/), followed by Kneaddata v.0.5.1 (--trimmomatic-options “LEADING:20 TRAILING:20 SLIDINGWINDOW:4:20 MINLEN:50”) for quality trimming, low-quality read removal and human reads decontamination. To identify *Desulfovibrio* MAGs, quality-controlled reads were assembled into contigs using Megahit v.3.14.1 with default parameters. MetaWRAP v.1.3.2 [111] was used for contig binning and refinement, using three binning methods (metabat2, maxbin2, and concoct) for contigs >1kb. The binning and bin annotation pipeline is published on https://github.com/GRONINGEN-MICROBIOME-CENTRE/GMH_MGS_pipeline.

### Phylogenetic analysis of *Desulfovibrio* genomes

Only genomes with completeness >90% and <5% contamination (estimated by CheckM [112]) were included in our analysis. All the *Desulfovibrio* genomes were de-replicated using dRep [113] with a 99.9% ANI cutoff. Genome taxonomy was assigned with GTDB-tk version 2.1.1 [41] and phylogeny of the *Desulfovibrio* genomes was constructed using Mashtree [114]. The visualization was done with GraPhlAn [115] based on Mashtree. To identify new species, we also calculated ANI and mash distance between *Desulfovibrio* genomes that can not be assigned to any existing species. An ANI value of >95% and mash distance <5% were used for defining species. ANI calculation was implemented using pyani [116] and mash distance was computed by Mash [117]. Genomes clustering was done using the neighbor-joining algorithm and visualized with the R package stats.

### Genome annotation and functional analysis of *Desulfovibrio* genomes

The genome annotation of MAGs, including coding sequence (CDS), tRNA and rRNA prediction, was performed with prokka [118]. All predicted CDS were annotated with multiple databases targeting different types of functions. Clusters of orthologous groups (COG) were annotated with EggNOG-mapper v2.1.64 [119] with default parameters. COG functional category frequencies were calculated as the proportion of genes involved in each category. InterProScan (v 5.59-91.0) [120] was used to predict functional domains for all proteins and AlphaFold2 [121] was used to predict flagellin gene structures based on amino acid sequences. Antimicrobial resistance genes and virulence factors were identified using ABRicate (https://github.com/tseemann/abricate) with the following settings: a minimum nucleotide identity of 60 % and a minimum DNA coverage of 80 %. Resfinder (for antimicrobial resistance) and VFDB (for virulence factor) annotation were used for downstream analysis.

### Pangenome analysis of *Desulfovibrio* species

Roary [122] was used to reconstruct the core- and pan-genome for each major *Desulfovibrio* species (i.e., containing >50 genomes) with minimum amino acid identity of 90% (‘-i 90’) and core genes defined at 99% presence (‘-cd 99’). A maximum likelihood phylogenetic tree was generated from the aligned sequences of core genes using FastTree v2.1.11. Permutation analysis of variance in gene content of *Desulfovibrio* species was performed on the matrix of pairwise Jaccard distances using the adonis2 function (R package Vegan) [123].

### Identification of genes involved in sulfur metabolism

A curated database of sulfur metabolism genes and their corresponding pathway information was constructed from the following resources: 1) KEGG database [65]; 2) Wolf et al. [66], 3) Braccia et al. [25]. Using a Hidden Markov Model (HMM), sulfur metabolism genes were subsequently searched against the predicted protein sequences of all *Desulfovibrio* genomes using hmmscan (v3.3.2, genes with e-values ≤ 1E-10 were counted as sulfur metabolism genes) [124]. In case of multiple hmm files corresponding to the same sulfur metabolic gene, the protein sequence was considered as a sulfur metabolic gene if the protein mapped to at least one of the hmm files.

### Identification of MAGs and sulfur metabolic genes in IBD

Stool samples from the cohort FedericiS_2022 [68] include a total of 15,002 Gbp of metagenomic quality-controlled reads from four populations (France, Germany, Israel, and the US) (range = 0.41–97.12 Gbp, avg = 27.94 Gbp, std dev = 25.16 Gbp). We identified 13,366 high-quality MAGs (≥90% completeness, ≤5% contamination) assigned to 16 phyla, 580 genera, and 1,528 species from 537 samples. Among these MAGs, 54 *Desulfovibrio* from 9 species and uncharacterized *Desulfovibrio spp.* were identified and present in 53 samples (**Supplementary Table 1**).

To identify sulfur metabolic genes at microbial community level, we generated a non-redundant gene catalog and profiled gene family abundances. Briefly, protein-coding genes were predicted using Prodigal [125, 126]. Then species-level non-redundant gene catalogs (identity >95%, coverage >90%) were generated for each of the four populations of this cohort (France, Germany, Israel, and the US) before applying cd-hit-est [127]. Afterwards, the representative sequences from all four populations were merged and grouped into the final gene catalog using the same thresholds. The representative sequences were then mapped against the hmm file of curated sulfur metabolism genes to identify relevant gene families at the community level (e-value = 1E-10). Finally, gene abundance was calculated by mapping the quality-controlled reads using bowtie2 [128] and normalizing as RPKM.

### Identification of tetrathionate metabolism related gene expression level in HMP2 metatranscriptomic data

Paired metagenomic and metatranscriptomic data from the HMP2 IBD cohort [79] was used to investigate gene activity related to tetrathionate metabolism. Quality controlled metagenomic and metatranscriptomic data was downloaded from https://ibdmdb.org/downloads/. Based on the metagenomic data, protein-coding genes were predicted using Prodigal [125] and a gene catalog was constructed with cd-hit-est with identity >95% and coverage >90% [127]. The expression of each gene family was calculated per sample by mapping the metatranscriptomic reads against the gene catalog using Bowtie2 (v2.4.1). Reads per kilobase per million mapped reads (RPKM) were calculated with SAMtools (v1.10).

### Antimicrobial susceptibility testing of *Desulfovibrio* isolates

*Desulfovibrio* isolates were grown in modified Postgate Medium for 48 h. 100 µl of each culture was spread thinly onto modified Gifu University Medical School (modified GAM, Himedia) Agar. Minimum Inhibitory Concentration (MIC) strips (Liofilchem) for Ampicillin, Ceftazidime, Chloramphenicol, Kanamycin, or Tetracycline were placed in the middle of the plate. Plates were incubated anaerobically (90 % N2, 5 % CO2, 5 % H2, 37 °C) for up to one week. Inhibition zones were measured after 48 hours and confirmed at the end of the incubation period. MIC was determined as the antibiotic concentration at the border of the inhibition zone.

### Swimming and swarming motility assays

*Desulfovibrio* isolates were grown in modified Postgate Medium for 48 h in biological triplicates. 10 µl of each culture was used to inoculate mGAM (Gifu University Medical School, Himedia) plates prepared with a 1.5 % agar base overlaid with 0.3 % or 0.6 % agar. Plates were incubated anaerobically (90 % N2, 5 % CO2, 5 % H2, 37 °C, 75 % humidity) for two weeks. Motility zones were measured daily, and the diameter of swarming or swimming expansion was recorded. The area of colony expansion was calculated using the formula A = π * a * b (https://rechneronline.de/pi/ellipse.php). Relative expansion was calculated using the area of colony expansion at the end of the incubation period compared to the area of colony expansion at the beginning of the incubation period.

### Detection of *Desulfovibrio* flagella by transmission electron microscopy (TEM)

*Desulfovibrio* strains were grown in liquid modified Postgate medium for 24 hours under anaerobic conditions (5% CO_2_, 5% H_2_, 90% N_2_) at 37 °C. 5 µl of the bacterial sample was pipetted onto a 200-mesh glow discharge treated continuous carbon copper grid and incubated for approximately 1 min. The samples were washed with 20 µl of buffer and negatively stained with 5 µl of a 2 % (w/v) uranyl acetate solution for 30 s. Excess solution was removed with filter paper. TEM micrographs were recorded on a JEOL JEM-1400 Plus transmission electron microscope at 120 kV with a Ruby (JEOL) CCD camera at a nominal magnification of 10,000 or 60,000. Final pixel size is 1.6 or 0.275 nm/px, respectively.

### TLR5 reporter cell line stimulation with bacterial culture supernatant filtrates

Frozen pure cultures (20% glycerol) of *Desulfovibrio* strains were revived on blood agar (Columbia agar plates, 7% sheep blood) under anaerobic conditions (5% CO_2_, 5% H_2_, 90% N_2_, 37 °C). Single colonies were incubated in 10 ml modified liquid Postgate medium (0.5g K2HPO4, 1.0g NH4Cl, 1.0g Na2SO4, 0.1g CaCl2 x 2 H2O, 2.0g MgSO4 x 7 H2O, 5.0g Sodium pyruvate, 1.0g Yeast extract, 0.5ml Na-resazurin (0.1% w/v), 0.1g Na-thioglycolate, 0.1g Ascorbic acid) for 48 hours in bacterial tubes. Bacterial suspensions were vortexed vigorously to allow detachment of flagella, centrifuged at 10 x g for 10 min, and sterile filtered to obtain supernatant. In parallel, HEK-Blue TLR5 reporter cell line (InvivoGen) reporting for murine TLR5 receptors and control cell line HEK-Blue Null1 without TLR gene overexpression (passages 7 to 9) were cultivated at 37 °C under 5% CO2 in Dulbecco MEM high glucose supplemented with 10% (v/v) fetal bovine serum, 10% (v/v) antibiotic antimycotic solution, 1mM L-Glutamine and specific antibiotics according to manufacturers’ recommendations. Cells were seeded in 48-wells cell culture plates according to their growth rate between cell lines to reach a density of 90% after 24h. Then, cells were washed with PBS and treated with sterile bacterial supernatant (diluted in cell culture medium without specific antibiotics 1:5, corresponding to 100 to 1000 bacteria per cell). Cells were incubated with treatment for 18h, then activation of TLR was assessed with the colorimetric enzyme assay QUANTI-Blue by InvivoGen according to manufacturer’s recommendations. The QUANTI-Blue Solution was left to incubate for 15 min, and activity was measured using a spectrophotometer at 620 nm.

### Recombinant flagellin expression and purification

Flagellins’ DNA was commercially cloned into pETM11-pLIB vector with 6x-histidine tag in N-terminal (Genwizz). Plasmid inserts were transformed into competent *E. coli* BL21(DE3) cells (NewEngland Biolabs). The transformed cells were grown at 37°C in LB supplemented with kanamycin (50 μg/ml) to an OD600 (optical density at 600 nm) of 0.4 to 0.6/ml. Protein expression was induced with 0.1 M isopropyl-β-D-thiogalactopyranoside (Sigma), and cells were incubated for 2.5 hours at 37°C. Cells were harvested by centrifugation (3400g for 15 min at 4°C), and pellets were resuspended in 15mL of lysis buffer [20 mM tris-HCl (pH 8), 300mM NaCl, 5mM Imidazole, 1% triton-100, pH=7.5] + Halt protease inhibitor cocktail (ThermoFisher Scientific). The lysate was sonicated four times with a 40% amplitude for 1 minute total followed by centrifugation (3400 g for 60 minutes at 4°C). The supernatant was filtered using 0.22uM filters.

For the flagellin purification, cleared lysates were incubated with Ni-NTA (nitrilotriacetic acid) agarose resin (ThermoFisher Scientific)(1:4 volume; QIAGEN) for 2 hours at room temperature. Agarose was then washed once with two column volumes (CVs) of wash buffer [20 mM tris-HCl, 300mM NaCl, 20mM Imidazole, pH=7.5]. Ni-NTA–bound proteins were eluted with 1 CV of lysis buffer). The eluted proteins were transferred in a 30kDa concentration tube (Sigma) and centrifuged (2000 g for 10 minutes at 4°C). The proteins were dialyzed at 4°C in PBS-Tween (0.01%) for 2 hours with buffer change every hour and overnight. Purified proteins were quantified by Qubit®, and aliquots were stored at −80°C.

### TLR5 reporter cell line assay using purified flagellins

For *in vitro* experiments with HEK-Blue mTLR5 and Null1-v reporter cell lines, the manufacturer’s instructions were followed for growth and antibiotic selection (Invivogen). For the TLR5 reporter assay, 20.000 cells/per well were seeded into a 96-well plate and treated with 0.5ng/ml of FlaBS and FlaST) and 1000 ng/ml of recombinant *Desulfovibrio* flagellins. The HEK-Blue Null1-v parental cell line was used as a background signal control. After 18 hours, the supernatant from each well was collected, mixed 1:1 with the HEK-Blue detection medium and incubated at 37°C in the dark for 90 minutes. Absorbance was measured at 630nm using a spectrophotometer.

### Small intestinal organoid culture

Small intestinal organoids were generated from isolated small intestinal crypts from 8 to 12-week-old C57BL/6 mice. They were embedded in 50% Matrigel (Corning Incorporated) domes and cultured in IntestiCult™ Organoid Growth Medium (Stemcell Technologies), supplemented with 1% sterile penicillin–streptomycin (Thermo Fisher Scientific). Organoids were passaged at least once before experiments were performed.

CRISPR-Cas9-mediated deletion of *Tlr5* in organoids was performed using the lentiCRISPRV2 lentiviral system (Addgene plasmid no. 52961) and a single-guide (sgRNA) targeting exon4 of *Tlr5* (target sequence AGGGAGATATTACCAACACG). Lentiviral particles were produced by HEK293T cells and transduced into organoids as previously described [62]. Following transduction, organoids were maintained under puromycin selection (4 µg/mL; InvivoGen) for two to three passages prior to further analysis. To assess the knock-out efficiency, the target site was PCR-amplified using the primers 5’-TACTGGTGCCCGTGTGTAAA-3’ and 5’-ACAGCCGAAGTTCCAAGAGA-3’, and verified by Sanger sequencing. Indel and knock-out scores were analyzed using ICE analysis (Inference of CRISPR Edits) from Synthego, yielding 91% and 62%, respectively [129].

### Small intestinal organoid treatment with purified flagellins

A low-viscosity medium consisting of IntestiCult™ OGM Mouse Basal Medium supplemented with 10% Matrigel™ was used for all experiments [130]. Wild-type and TLR5KO organoids were seeded into 96-well plates at a density of 100 −150 organoids per well in 100µl medium. After two days of growth (37°C, 5% CO2), the medium was aspirated and replaced with fresh medium supplemented with either of the following: 10nM flagellin from *B. subtilis* (FlA-BS; Invivogen) or *S.typhimurium* (FLA-ST; Invivogen), and 30nM recombinant *Desulfovibrio* flagellins. After 24h, organoids were dislodged by pipetting after incubating for 10 minutes on ice with 100µl of cold PBS containing 2 mM EDTA. The organoids were stored in RLT buffer from the RNeasy Minikit (QIAGEN) at −80°C until processing for RNA extraction per the manufacturer’s instructions (Macherey-Nagel). cDNA was synthesised from 500 µg of RNA using the SensiFAST cDNA synthesis kit (Meridian Bioscience) per manufacturer’s instructions and diluted 10-fold before use. qPCR was performed using the GoTaq qPCR master mix (Promega) on a LightCycler® 480 II (Roche). qPCR primer sequences were obtained from Primer Bank [131].

### Organoid Profile Analysis

Brightfield images acquired with a wide-field Leica THUNDER microscope were analyzed using the Napari Organoid Analyzer (NOA) [132] to extract morphological features, including roundness and perimeter. Roundness and perimeter were computed from the resulting masks and used with predefined thresholds to assign each organoid to enterosphere (perimeter 200–400 and roundness ≥ 0.85), budding (roundness < 0.85 and perimeter ≤ 400), or enteroid (roundness < 0.85 and perimeter > 400) stages across all treatment conditions.

### Bulk RNA sequencing of flagellin-treated organoid experiments

Messenger RNA was purified from total RNA using poly-T oligo-attached magnetic beads (Novogene). After fragmentation, the first strand cDNA was synthesized using random hexamer primers, followed by the second strand cDNA synthesis using either dTTP for non strand specific library or dUTP for strand specific library. For the non strand specific library, it was ready after end repair, A-tailing, adapter ligation, size selection, amplification, and purification. The library was checked with Qubit and real-time PCR for quantification and a bioanalyzer for size distribution detection. Quantified libraries were sequenced on Illumina platforms, according to effective library concentration and data amount.

Original image data file from the Illumina platform was transformed to sequenced reads (Raw Data) by CASAVA base recognition (Base Calling). Raw data was filtered to remove adapter contamination, reads with uncertain nucleotides constituting more than 10% of either read (N > 10%), and reads with low-quality nucleotides (Base Quality less than 5) constituting more than 50 % of the read. Sequence alignments were performed with HISAT2 with the Genome assembly GRCm39 (mm39) as the reference genome [133].

Read counts were imported into the DESeq2 R package, and differential gene expression analysis was performed using the standard pipeline and default settings [134]. Heatmaps of differentially expressed genes were produced with the pheatmap R package. Gene ontology enrichment analysis was performed using the ClusterProfiler [135], org.Mm.eg.db, and enrichplot R packages.

## Data availability

The *Desulfovibrio* genome database, including newly sequenced isolate assemblies and identified MAGs, are available at Zenodo: https://doi.org/10.5281/zenodo.17927643. The corresponding metadata information is included in **Supplementary Table 1**. Detailed information for all isolates can be found in **Supplementary Table 5**. All newly isolated strains have been characterized and were deposited at the Weihenstephaner Stammsammlung at the Core Facility Microbiome, Technical University of Munich with WDCM number 1163 (https://ccinfo.wdcm.org/details?regnum=1163) under accession number WS 5745 - WS 5752 (further details in **Supplementary Table 5**). For strain MS028 biobank deposition was not possible due to ethics restriction for EEN patient samples. Two strains (BP2_2_G5_PP-3-Pallet_E04 and BP2_2_G7_PP-1-Pallet_F11) were isolated and are available through openBiome [98]. The description of the new species has been added to **Supplementary information**.

