## Supplementary information for "Extensive genomic diversity in *Desulfovibrio* species reveals species-specific functional traits associated with disease"

**This file includes:**

**Supplementary Methods**

**Supplementary Figures 1-19**

**Supplementary Tables 1-11 [Separate Files]**

### **Supplementary Methods**

#### **Description of novel *Desulfovibrio* species**

##### **Description of *Desulfovibrio longus* sp. nov.**

*Desulfovibrio longus* sp. nov. (L. masc. adj. longus, long; to reflect the elongated cell morphology of the bacterium observed under transmission electron microscopy). The description of this species is based on the features of *Desulfovibrio sp900556755* isolates (BP2_1_F7_FSA-079_D05, BP2_1_F8_FSA-041_F10 and BP2_2_D2_FSA-076_A04). Species separation was confirmed by GTDB-Tk assignment of the strains annotated as “*Desulfovibrio sp900556755*”. Transmission electron microscopy showed that the cells are vibrio-shaped, elongated rods measuring approximately 2–6 µm in length; non-flagellated (confirmed by gene analysis). The genomic analysis showed that this species encodes chloramphenicol resistance gene catB7. API 20A testing of all three isolates showed no acid production from glucose, mannitol, lactose, saccharose, maltose, salicin, xylose, arabinose, glycerol, cellobiose, mannose, melezitose, raffinose, sorbitol, rhamnose or trehalose. The isolates were also negative for gelatin hydrolysis (protease) and esculin hydrolysis (β-glucosidase). KEGG-based analysis identified the presence of the following pathways: acetate production from acetyl-CoA (EC:2.3.1.8, 2.7.2.1), propionate production from propanoyl-CoA (EC:2.3.1.8, 2.7.2.1), L-glutamate production from ammonia via L-glutamine (EC:6.3.1.2, 1.4.1.4), sulfide and L-serine utilised to produce L-cysteine and acetate (EC:2.3.1.30, 2.5.1.47) and riboflavin (vitamin B2) biosynthesis from GTP (EC:3.5.4.25, 3.5.4.26, 4.1.99.12, 2.5.1.78, 2.5.1.9, 2.7.1.26). The G+C content of genomic DNA is 63.24% - 63.51%.

##### **Description of *Desulfovibrio groningensis* sp. nov.**

*Desulfovibrio groningensis* sp. nov. (N.L. masc. adj. groningensis, of Groningen, the site of isolation). The description of this species is based on the features of the isolate BP-2_FSA-089_A08. Species separation was confirmed by GTDB-Tk assignment of the isolate as “uncharacterized *Desulfovibrio* spp.”. Transmission electron microscopy showed that the cells are vibrio-shaped, elongated rods measuring approximately 1–2 µm in length. The strain showed medium swimming and swarming activity compared to other *Desulfovibrio* species (expansion < 300%). The genomic analysis identified the presence of flagellar genes in this isolate. On mGAM, this isolate exhibited Chloramphenicol resistance. API 20A testing of the isolate showed no acid production from glucose, mannitol, lactose, saccharose, maltose, salicin, xylose, arabinose, glycerol, cellobiose, mannose, melezitose, raffinose, sorbitol, rhamnose, or trehalose. The isolate was also negative for gelatin hydrolysis (protease) and esculin hydrolysis (β-glucosidase). KEGG-based analysis identified the presence of the following pathways: acetate production from acetyl-CoA (EC:2.3.1.8, 2.7.2.1), propionate production from propanoyl-CoA (EC:2.3.1.8, 2.7.2.1), L-glutamate production from ammonia via L-glutamine (EC:6.3.1.2, 1.4.1.4), sulfide and L-serine utilised to produce L-cysteine and acetate (EC:2.3.1.30, 2.5.1.47) and riboflavin (vitamin B2) biosynthesis from GTP (EC:3.5.4.25, 3.5.4.26, 4.1.99.12, 2.5.1.78, 2.5.1.9, 2.7.1.26). The G+C content of genomic DNA of the isolate is 57.49%.

##### **Description of *Desulfovibrio* *paradesulfuricans* sp. nov.**

*Desulfovibrio* *paradesulfuricans* sp. nov. (Gr. prep. para, next to, resembling; L. prep. de, from; L. neut. n. sulfur, sulfur (S); N.L. masc. part. adj. desulfuricans, reducing sulfur compounds; from N.L. v. desulfurico, to reduce sulfur; denoting the closeness to *D. desulfuricans*). The description of this species is based on the features of the two isolates BP-3_FSA-069_C06 and BP-2_FSA-147_E02. Species separation was confirmed by GTDB-Tk assignment of the strains annotated as “*Desulfovibrio desulfuricans_A*”. Both isolates showed medium swimming and swarming activity compared to other *Desulfovibrio* species (expansion < 300%). The genomic analysis identified the presence of flagellar genes in the two isolates assigned to this species. On mGAM, the two isolates exhibited kanamycin resistance. API 20A testing of the isolates showed no acid production from glucose, mannitol, lactose, saccharose, maltose, salicin, xylose, arabinose, glycerol, cellobiose, mannose, melezitose, raffinose, sorbitol, rhamnose, or trehalose. The isolates were also negative for gelatin hydrolysis (protease) and esculin hydrolysis (β-glucosidase). KEGG-based analysis identified the presence of the following pathways: acetate production from acetyl-CoA (EC:2.3.1.8, 2.7.2.1), propionate production from propanoyl-CoA (EC:2.3.1.8, 2.7.2.1), L-glutamate production from ammonia via L-glutamine (EC:6.3.1.2, 1.4.1.4), sulfide and L-serine utilised to produce L-cysteine and acetate (EC:2.3.1.30, 2.5.1.47) and riboflavin (vitamin B2) biosynthesis from GTP (EC:3.5.4.25, 3.5.4.26, 4.1.99.12, 2.5.1.78, 2.5.1.9, 2.7.1.26). The G + C content of genomic DNA is 57.83% - 58.2%.

### **Supplementary Figures**

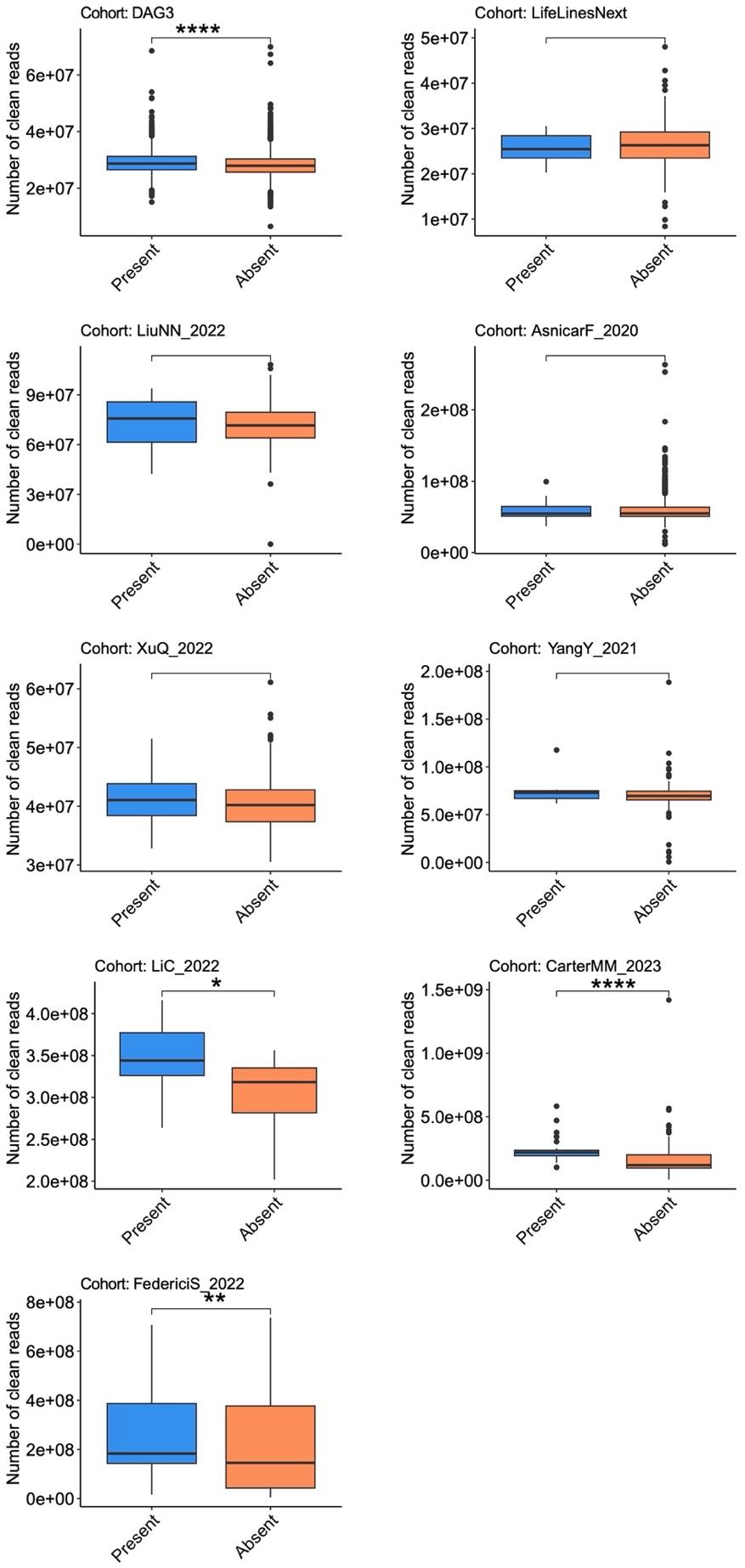

**Supplementary Fig. 1. Sequencing depth comparison of samples with and without detected *Desulforvibrio* metagenome assembled genomes (MAGs).** Sequencing depth represents the number of quality-controlled metagenomic reads. Cohorts used for de novo MAG recovery with at least 10 *Desulfovibrio* MAGs were included. Wilcoxon test: * p < 0.05; ** p<0.01; *** p<0.001; **** p<0.0001.

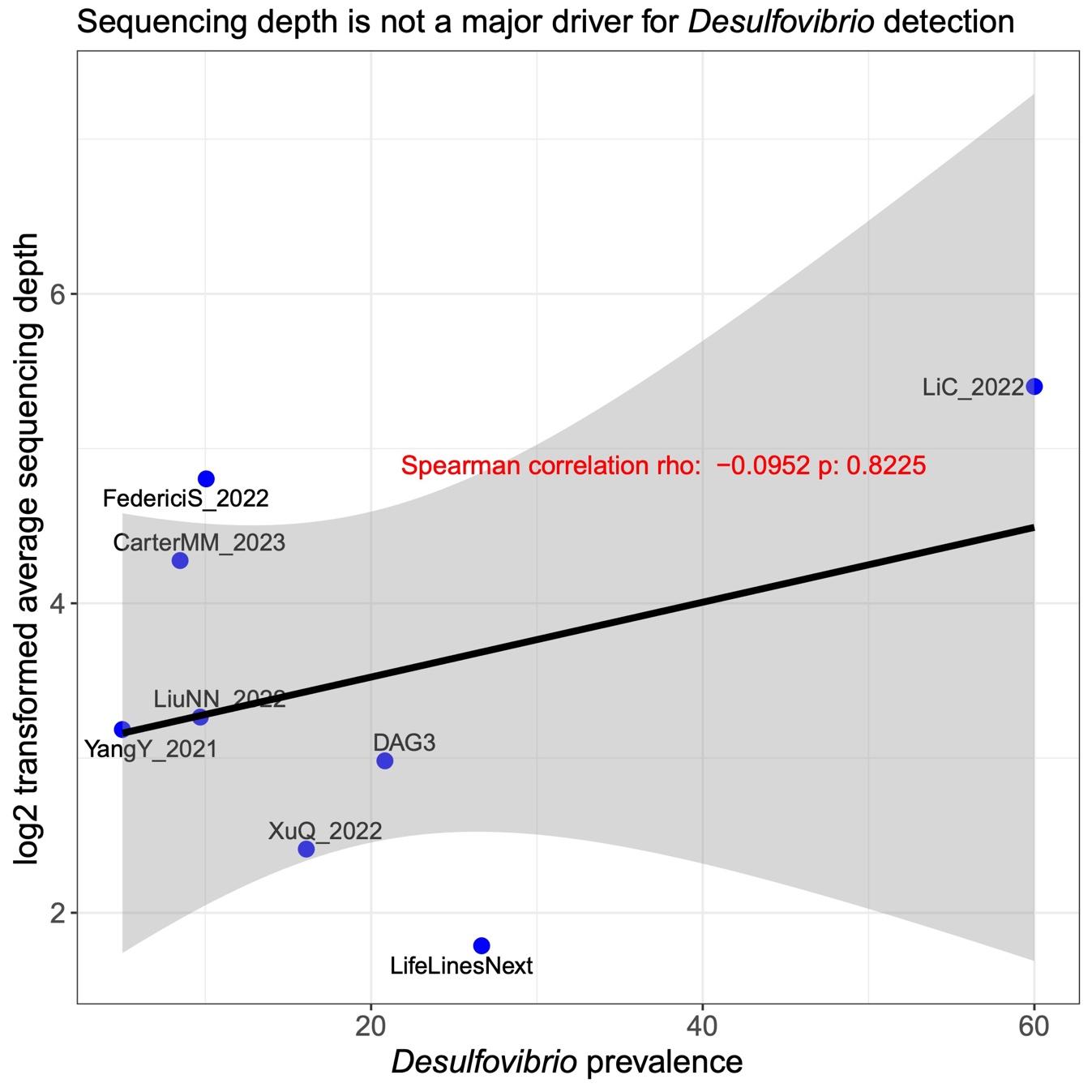

**Supplementary Fig. 2. *Desulfovibrio* prevalence versus sequencing depth across different cohorts.** Sequencing depth was measured as the number of total bases of quality-controlled metagenomic reads. Each point represents a cohort, where only cohorts with at least 10 Desulfovibrio MAGs were considered.

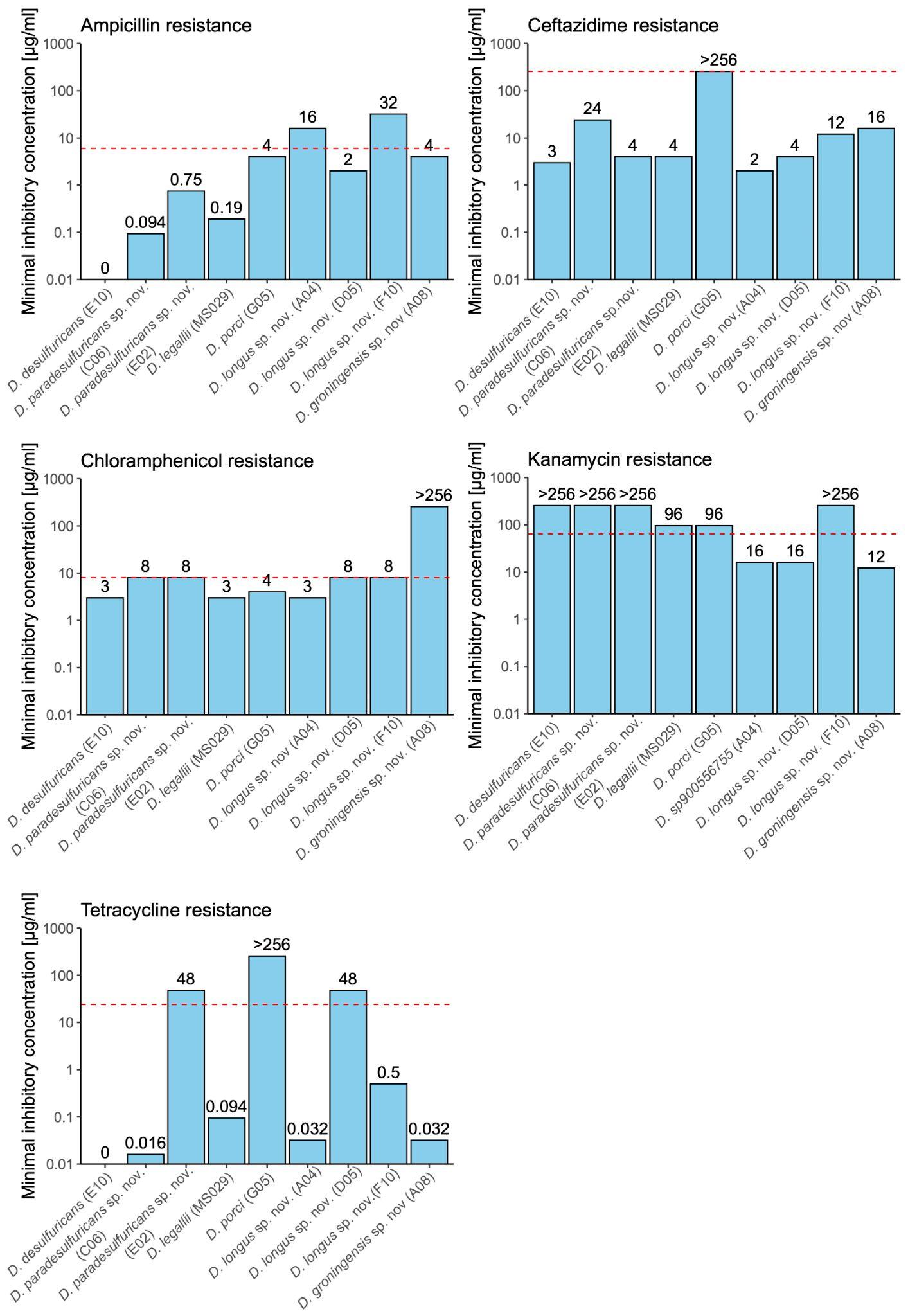

**Supplementary Fig. 3. *In vitro* antimicrobial susceptibility testing of *Desulfovibrio* isolates.** Dashed horizontal lines indicate the minimum inhibitory concentrations (MIC) clinical breakpoint used in this study. Abbreviations of the *Desulfovibrio* isolates on the x-axis are defined as follows: E10, BP-1_FSA-093_E10; C06, BP-3_FSA-069_C06; E02, BP-2_FSA-147_E02; G05, BP-3_FSA-069_G05; A04, BP2_2_D2_FSA-076_A04; D05, BP2_1_F7_FSA-079_D05; F10, BP2_1_F8_FSA-041_F10; A08, BP-2_FSA-089_A08.

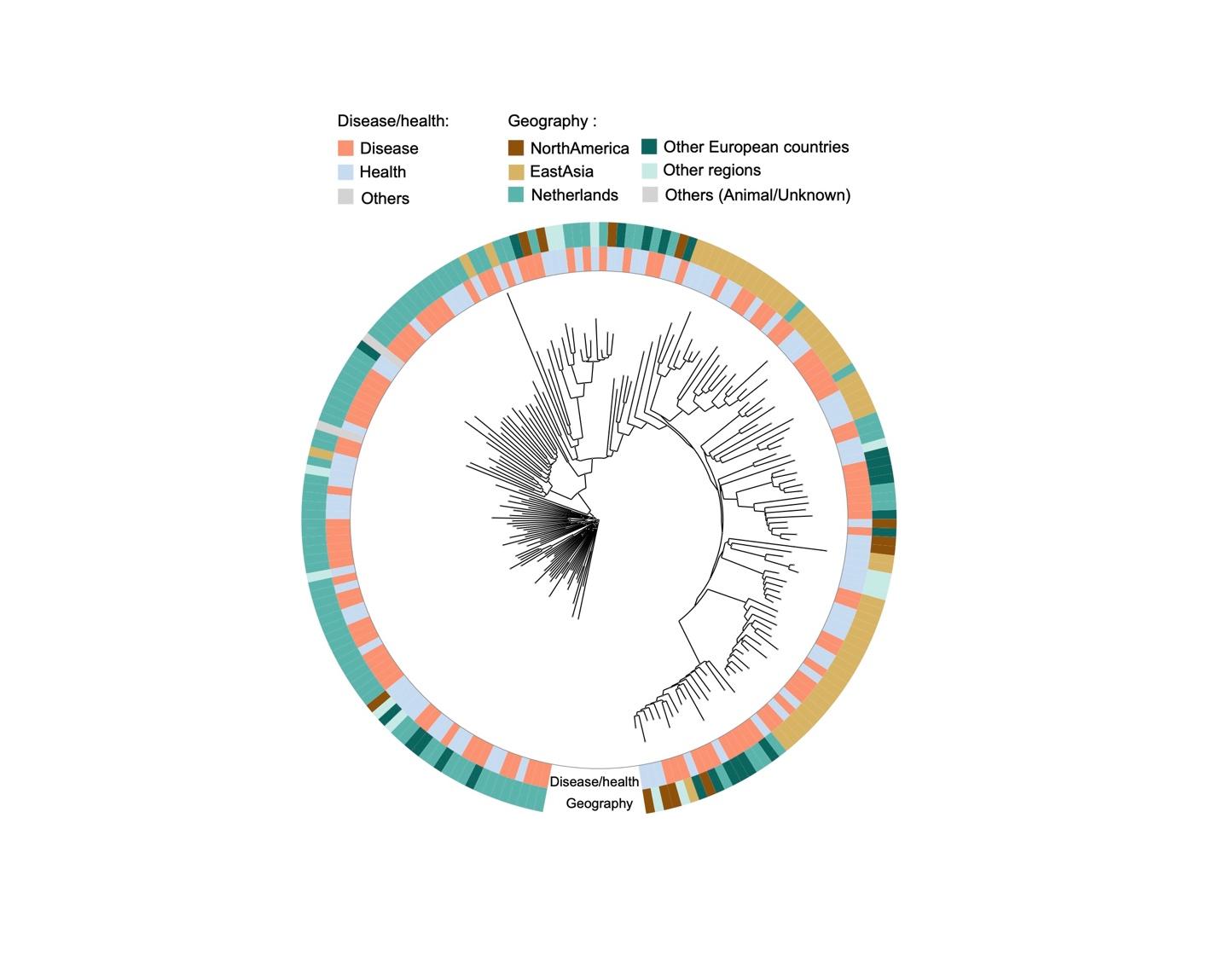

**Supplementary Fig. 4. Phylogenetic tree of *Desulfovibrio sp900319575* based on mash distance.** The rings indicate host phenotype (disease *vs.* health, “Others” refers to the genomes from animal samples or unknown sources) and host geography.

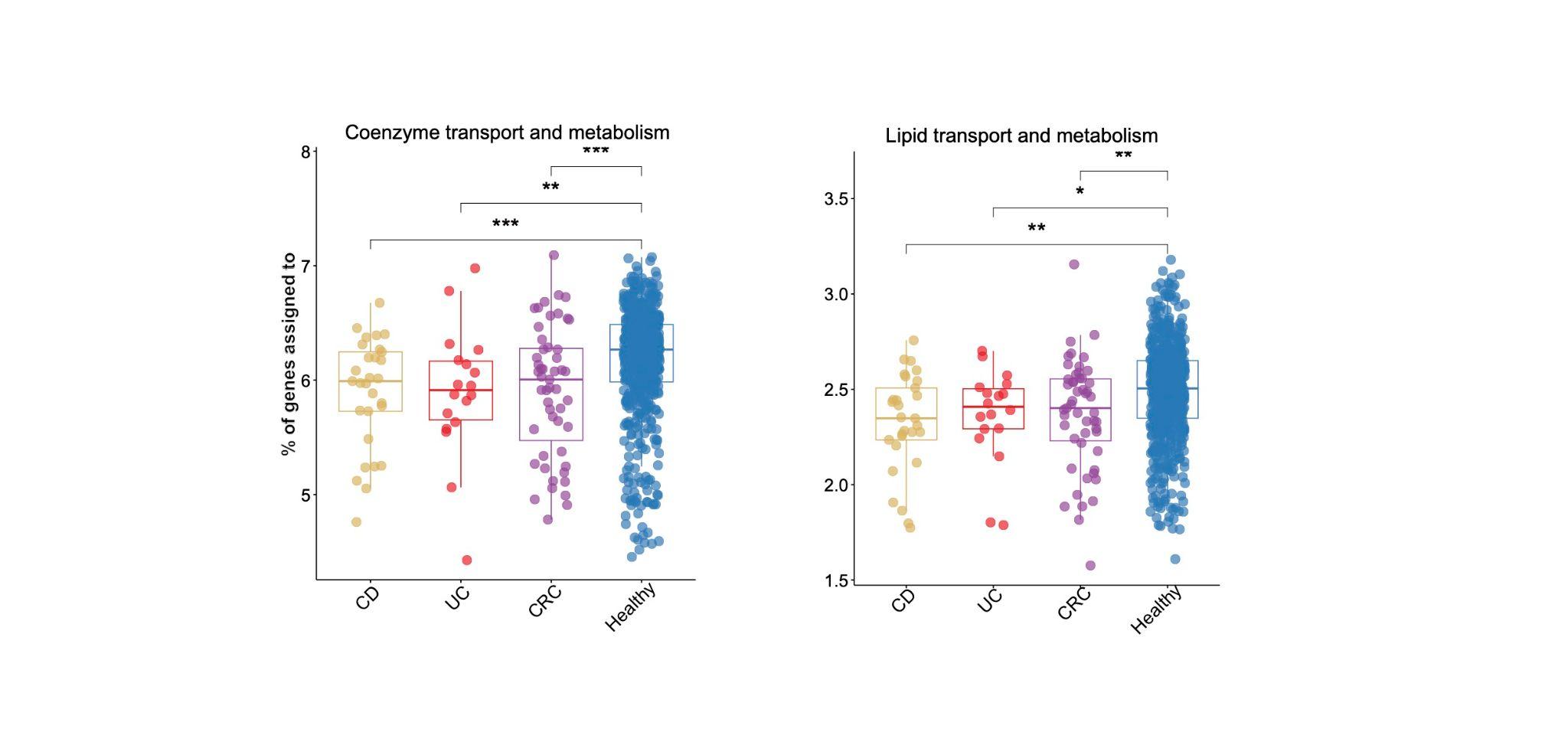

**Supplementary Fig. 5. Boxplots illustrate two broad clusters of orthologous groups (COG) that are depleted in *Desulfovibrio* spp. from inflammatory bowel diseases (IBD) and colorectal cancer (CRC) patients compared to healthy individuals** (UC, CD and CRC *vs.* healthy, Wilcoxon test, p<0.05). Stars represent different significance level: * p < 0.05; ** p<0.01; *** p<0.001**.**

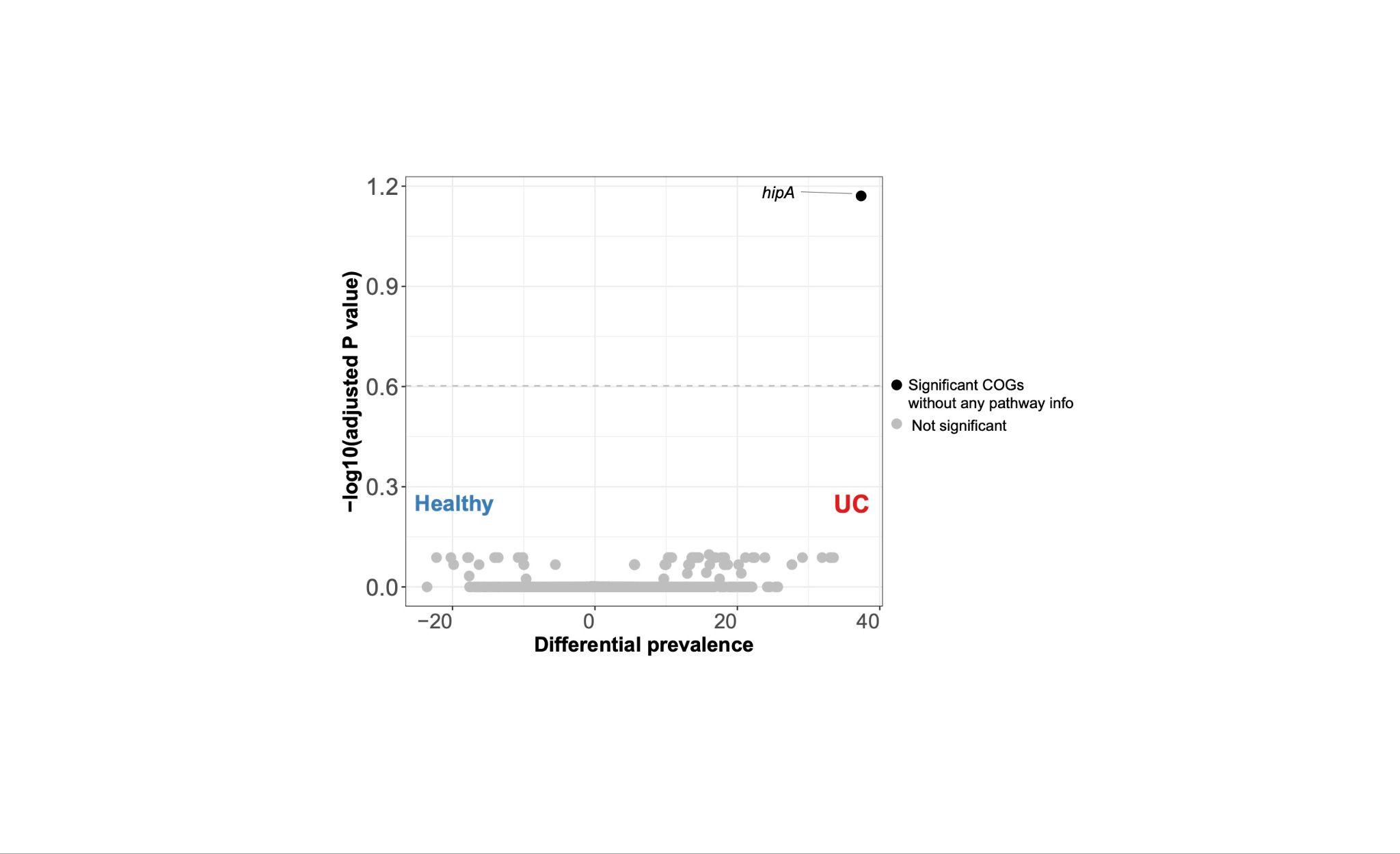

**Supplementary Fig. 6. Volcano plots of Clusters of Orthologous Genes (COGs) that are differentially enriched in ulcerative colitis (UC) compared to healthy individuals.** The x-axis indicates average COG prevalence in UC patients minus average COG prevalence in healthy individuals. Y-axis shows log-transformed adjusted p-value (Fisher’s exact test with Benjamini-Hochberg correction for multiple testing). The dashed horizontal line indicates the significance threshold (adjusted p-value = 0.25).

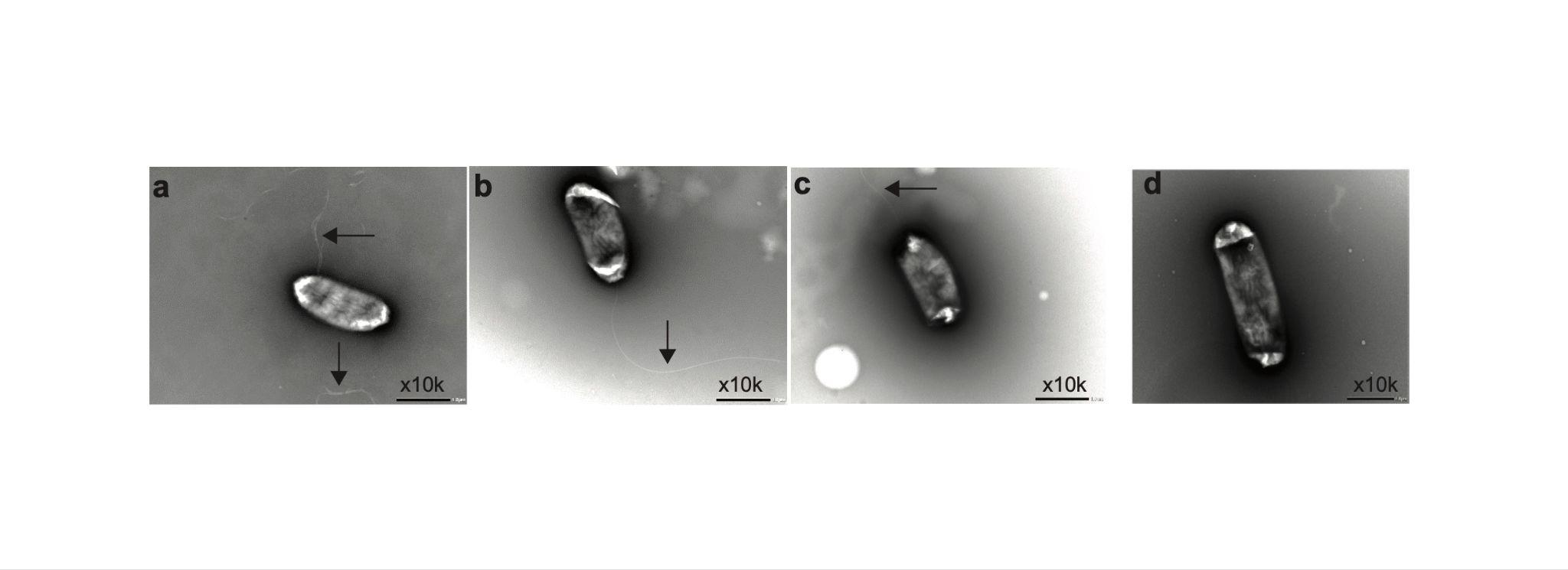

**Supplementary Fig. 7. Transmission electron microscopy of *D. desulfuricans*, *D. fairfieldensis*, *D. legallii* and *D. longus* sp. nov.** (**a**) *Desulfovibrio desulfuricans* (MS021) with visible but diffused flagella. (**b**) *Desulfovibrio fairfieldensis* (MS026) and (**c**) *Desulfovibrio legallii* (MS029) with visible flagella, but *D. sp900556755* (MS012) has no visible flagella.

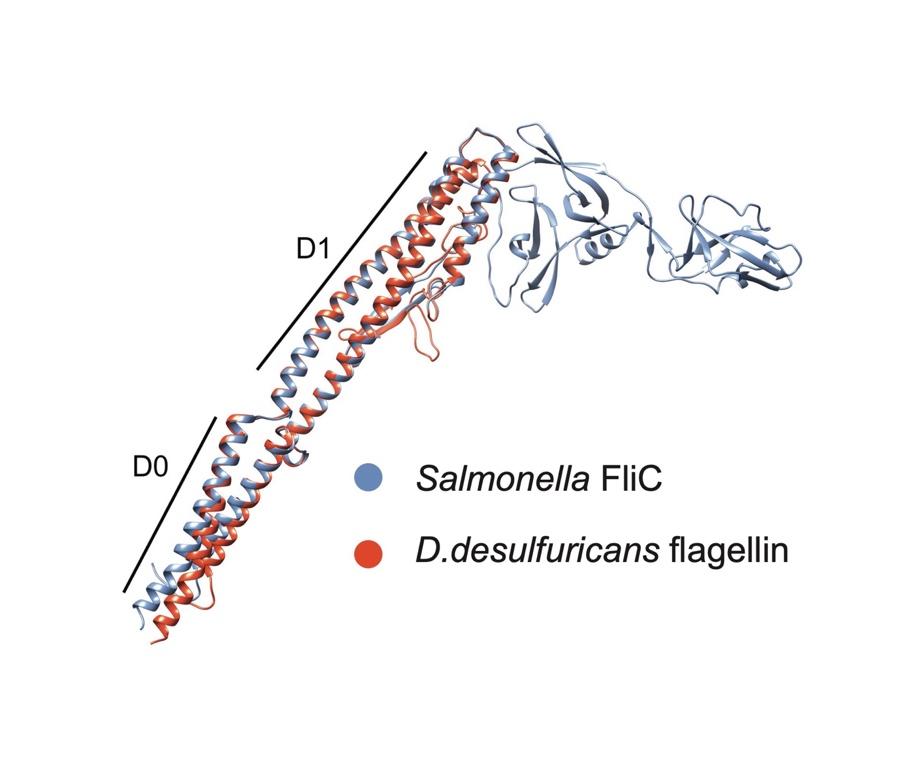

**Supplementary Fig. 8. Comparison of the structures of the flagellin of *Desulfovibrio desulfuricans* (MS020) and *Salmonella* FliC.** The structure of *D. desulfuricans* flagellin (red) is predicted by AlphaFold2, and the structure of Salmonella FliC (blue) is retrieved using PDB 1UCU. Chimera was used for the visualization.

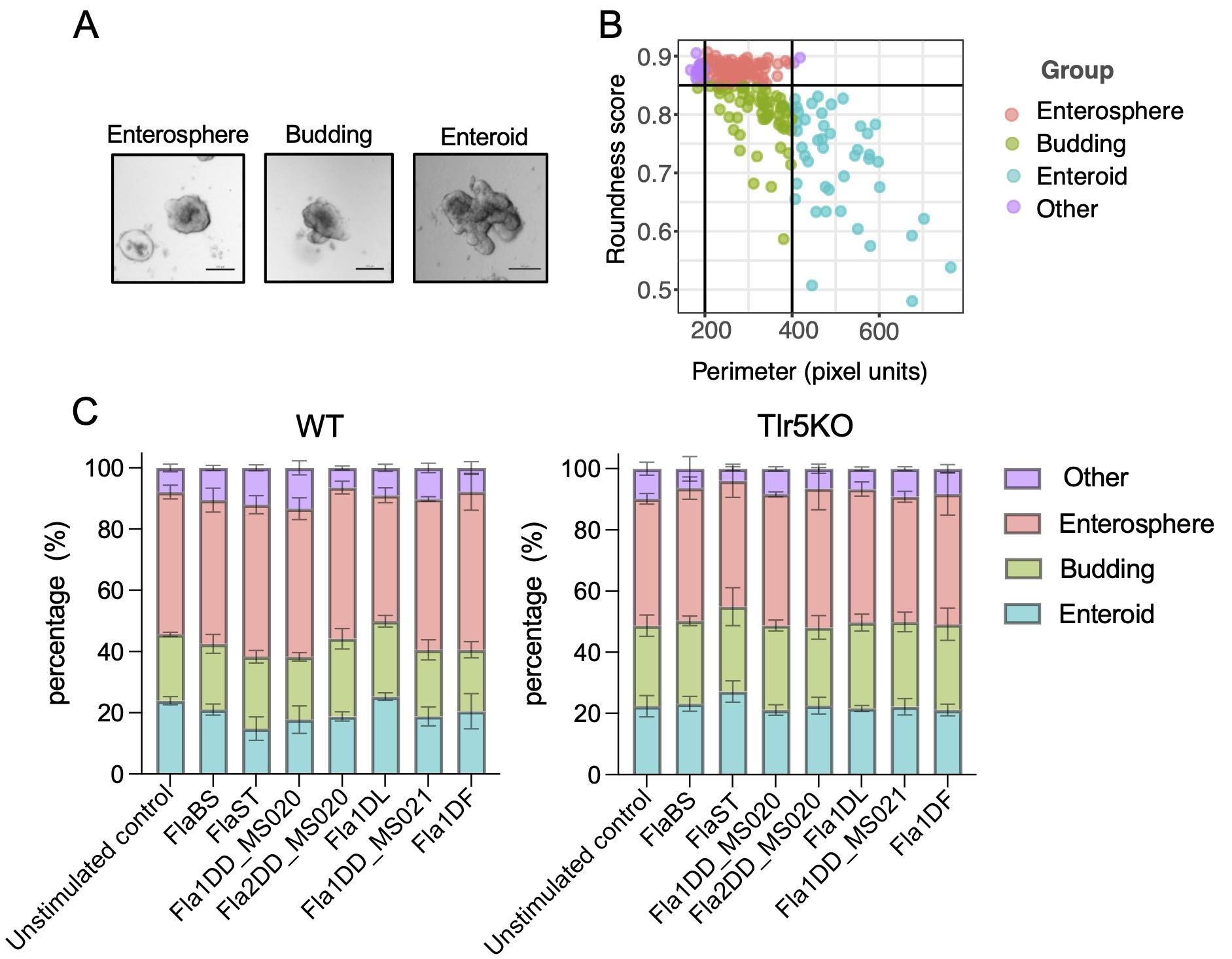

**Supplementary Fig. 9. Organoid profiling based on morphological features in response to different flagellin proteins.** (A) Representative stages of organoid development. Scale bar: 100 µm. (B) Gating strategy for classifying organoids based on morphology features extracted using NOA. Organoids were categorized as enterospheres (200 ≤ perimeter ≤ 400, roundness ≥0.85), budding organoids (roundness <0.85, perimeter ≤400), or enteroids (roundness <0.85, perimeter >400). (C) Distribution of organoid developmental stages across different treatments. Images from three biological replicates per condition were analyzed. All organoids within a single well were included in the analysis, with approximately 50–300 organoids quantified per well. Abbreviations of the flagellins on the x-axis are defined as follows: FlaBS, flagellin from *Bacillus subtilis;* FlaST: flagellin from *Salmonella typhimurium*; Fla1DD_MS020, flagellin 1 from *D. desulfuricans* (isolate MS020); Fla2DD_MS020, flagellin 2 from *D. desulfuricans* (isolate MS020); Fla1DD_MS021, flagellin 1 from *D. desulfuricans* (isolate MS021); Fla1DF: flagellin 1 from *D. fairfieldensis* (isolate MS026); Fla2DF: flagellin 2 from *D. fairfieldensis* (isolate MS026); Fla1DL: flagellin 1 from *D. legallii* (isolate MS029).

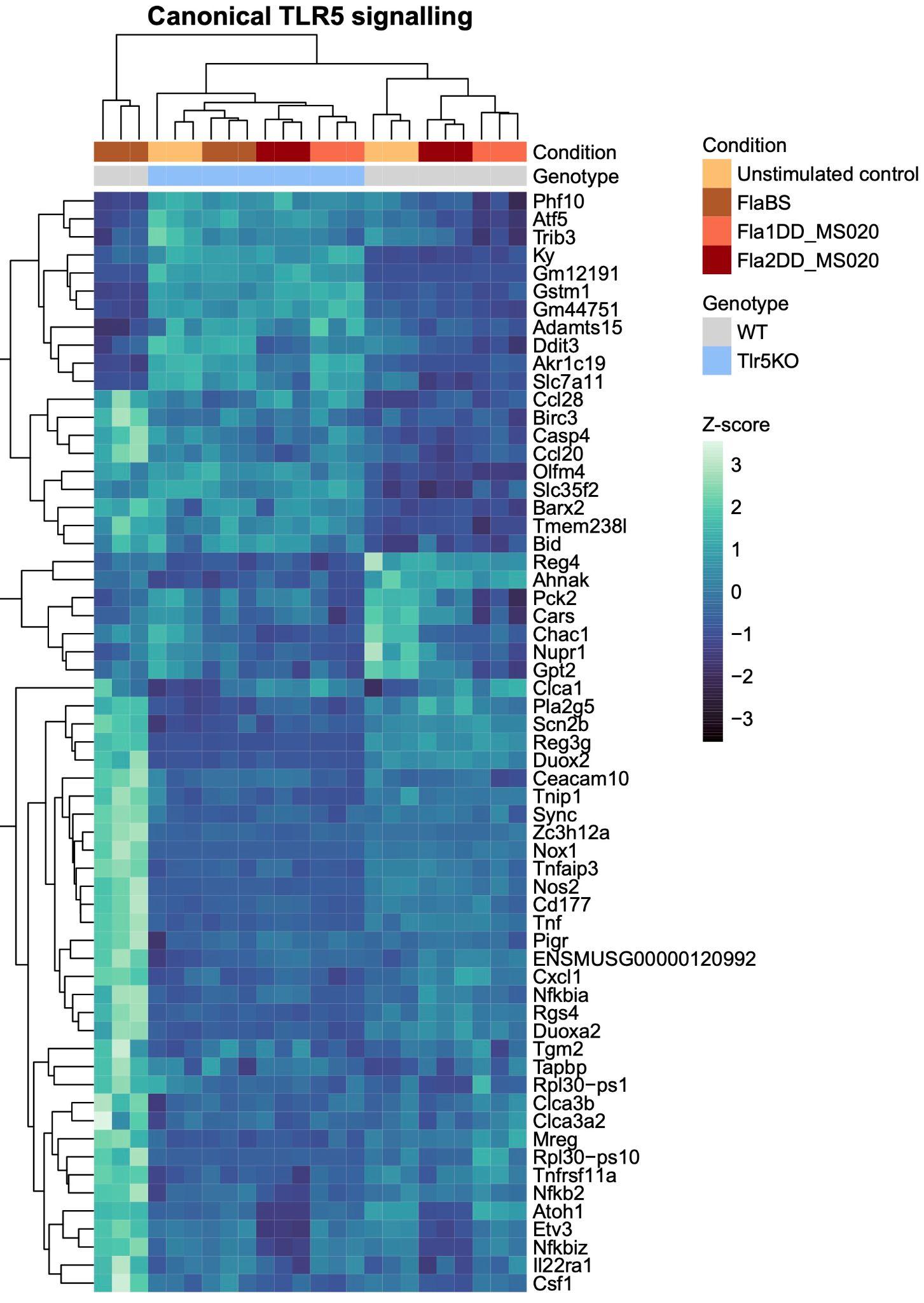

**Supplementary Fig. 10. Expression levels of canonical TLR5 signatures in wildtype and TLR5 knockout mouse organoids.** While FlaBS increased the expression of TLR5 signature including TNF, NF-κB, and antimicrobial peptide genes as expected as compared to unstimulated control (Wald test implemented in R package DESeq2, adjusted p <0.05), Fla1DD_MS020 and Fla2DD_MS020 did not induce the significant change of TLR5 signature.

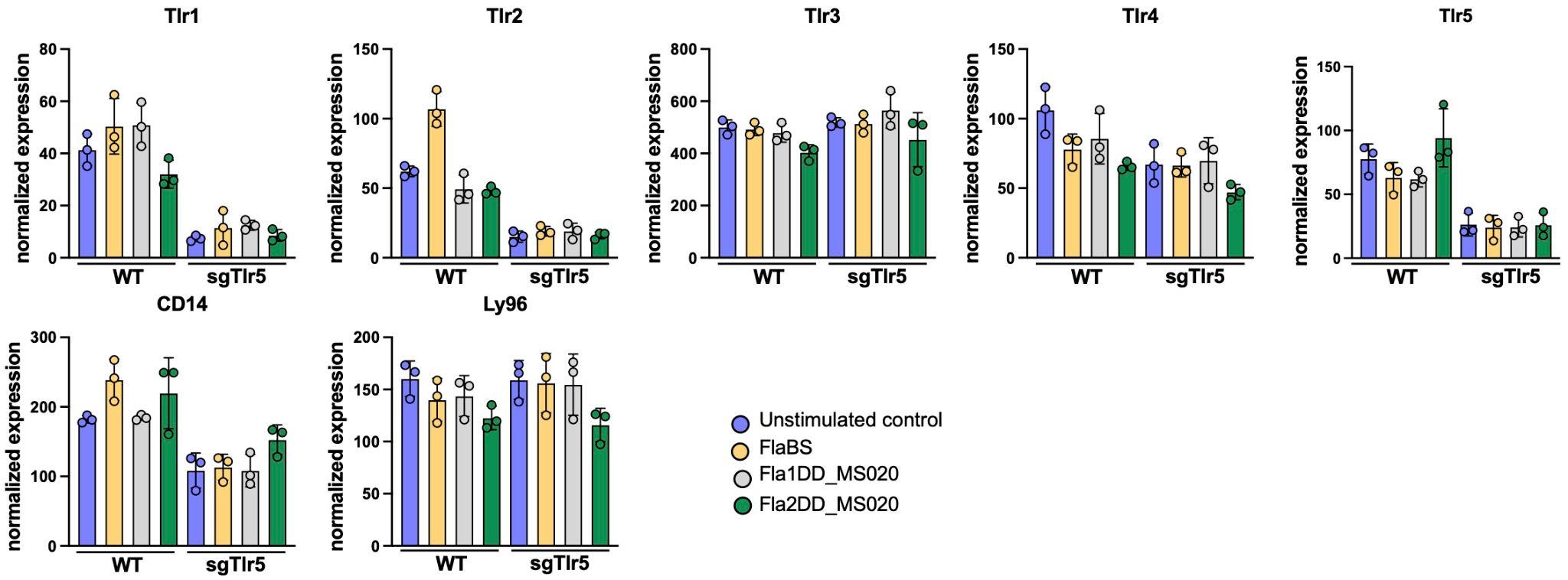

**Supplementary Fig. 11. Expression levels of TLRs regulated by *D. desulfuricans* flagellins in wildtype and TLR5 knockout mouse organoids.**

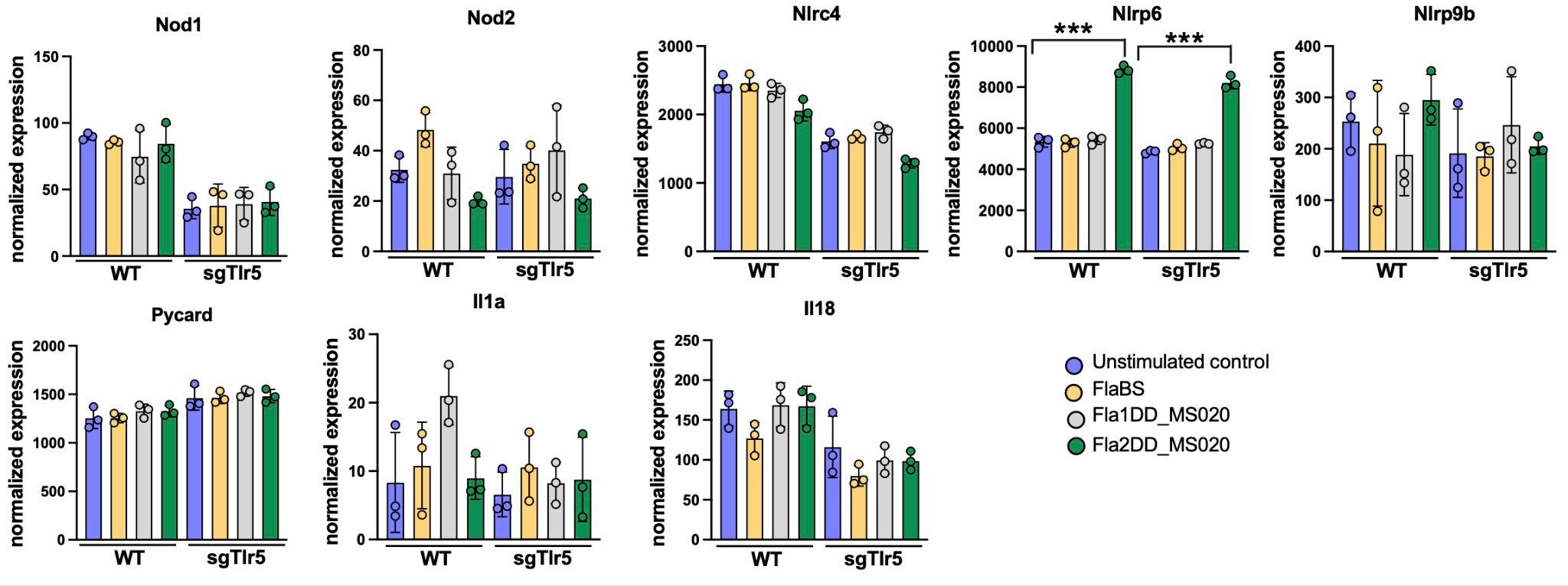

**Supplementary Fig. 12. Expression levels of the NLR family regulated by *Desulfovibrio desulfuricans* flagellins in wildtype and TLR5 knockout mouse organoids (Wald test).** Only *Nlrp6* showed strong upregulation by Fla2DD_MS020. Stars represent different significance level: *** p<0.001.

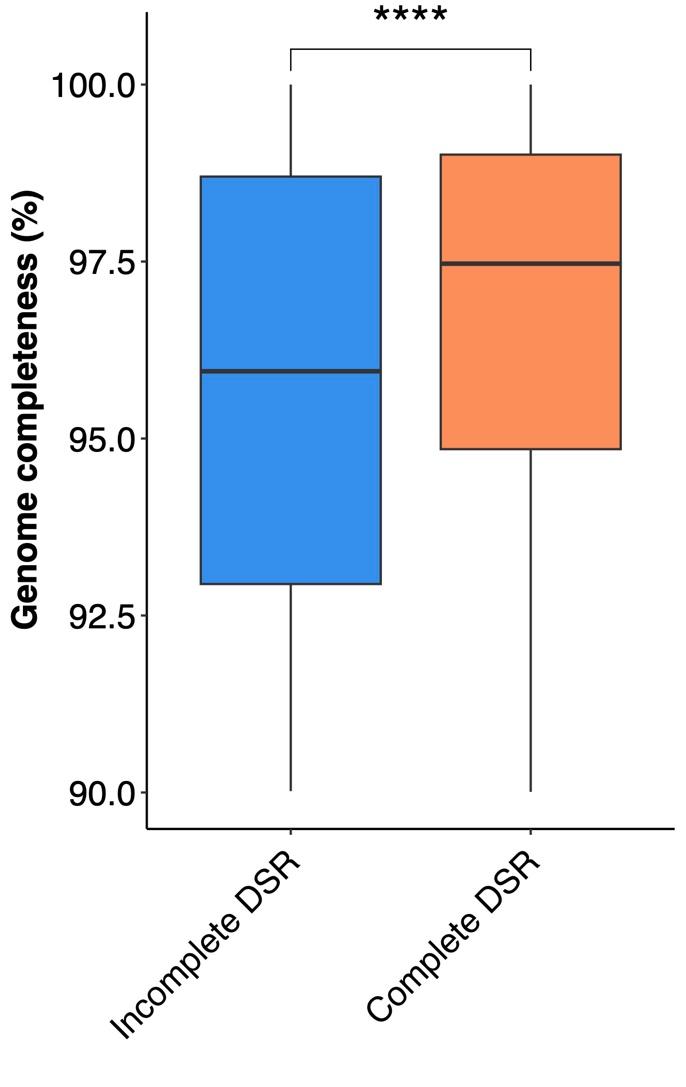

**Supplementary Fig. 13. Incomplete encoding of the dissimilatory sulfate reduction (DSR) pathway in *Desulfovibrio* genomes was associated with lower genome completeness (Wilcoxon test, p-value=8.617e-11).**

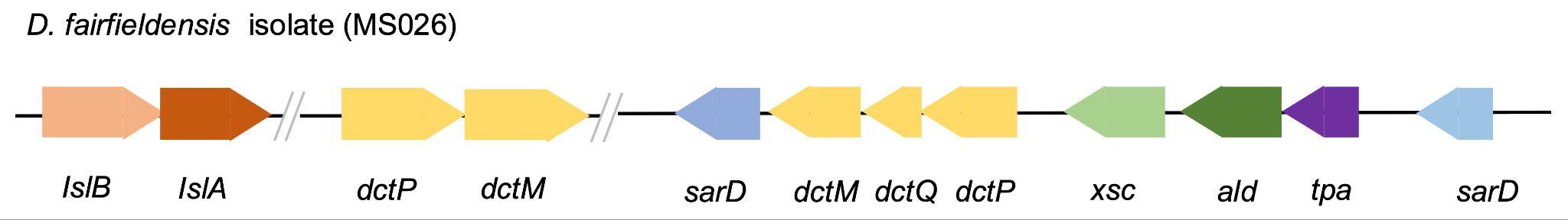

**Supplementary Fig. 14. Taurine metabolism gene clusters in a *D. fairfieldensis* isolate.** *D. fairfieldensis* encodes *dctPQM* genes that belong to tripartite ATP-independent periplasmic transporters (TRAP transporters, indicated in yellow), which facilitate the uptake of taurine from the environment, while other genes in the cluster (*tpa*, *xsc*, *ald* and *sarD*) complete the process of degrading taurine to sulfite. Specifically, taurine–pyruvate aminotransferase (*tpa*) converts taurine to sulfoacetaldehyde, which is further processed by sulfoacetaldehyde acetyltrasferase (*xsc*) to produce sulfite and *sarD* supports this process through the metabolism of sulfonate-derived aldehydes. Alanine dehydrogenase (*ald*) turns the resulting intermediate (L-Alanine) to pyruvate.

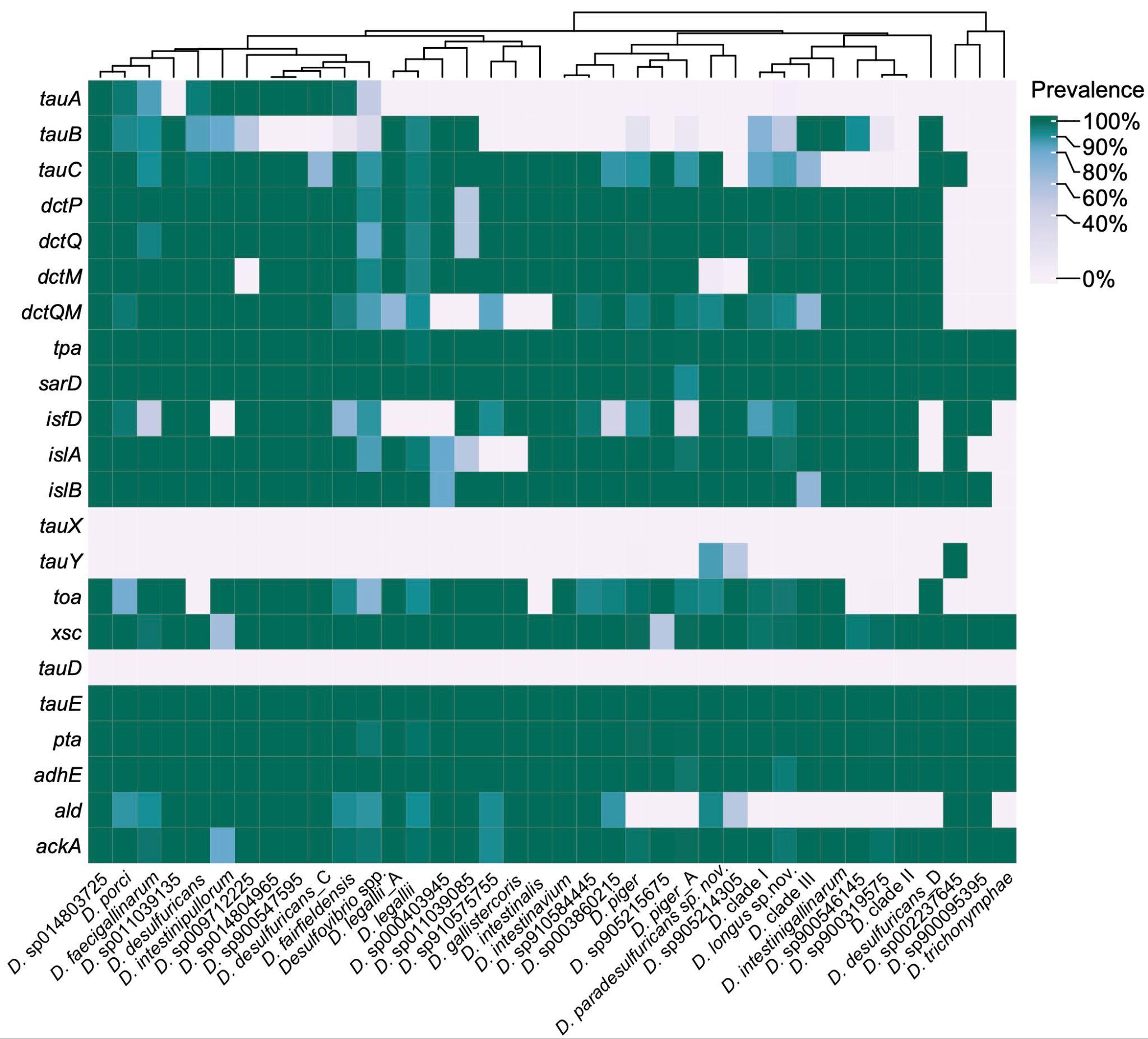

**Supplementary Fig. 15. Heatmap showing the presence of genes related to taurine metabolism in *Desulfovibrio* species.** The taurine metabolism genes were searched in all *Desulfovibrio* genomes (hmmscan, gene hits with e-value≤1E-10).

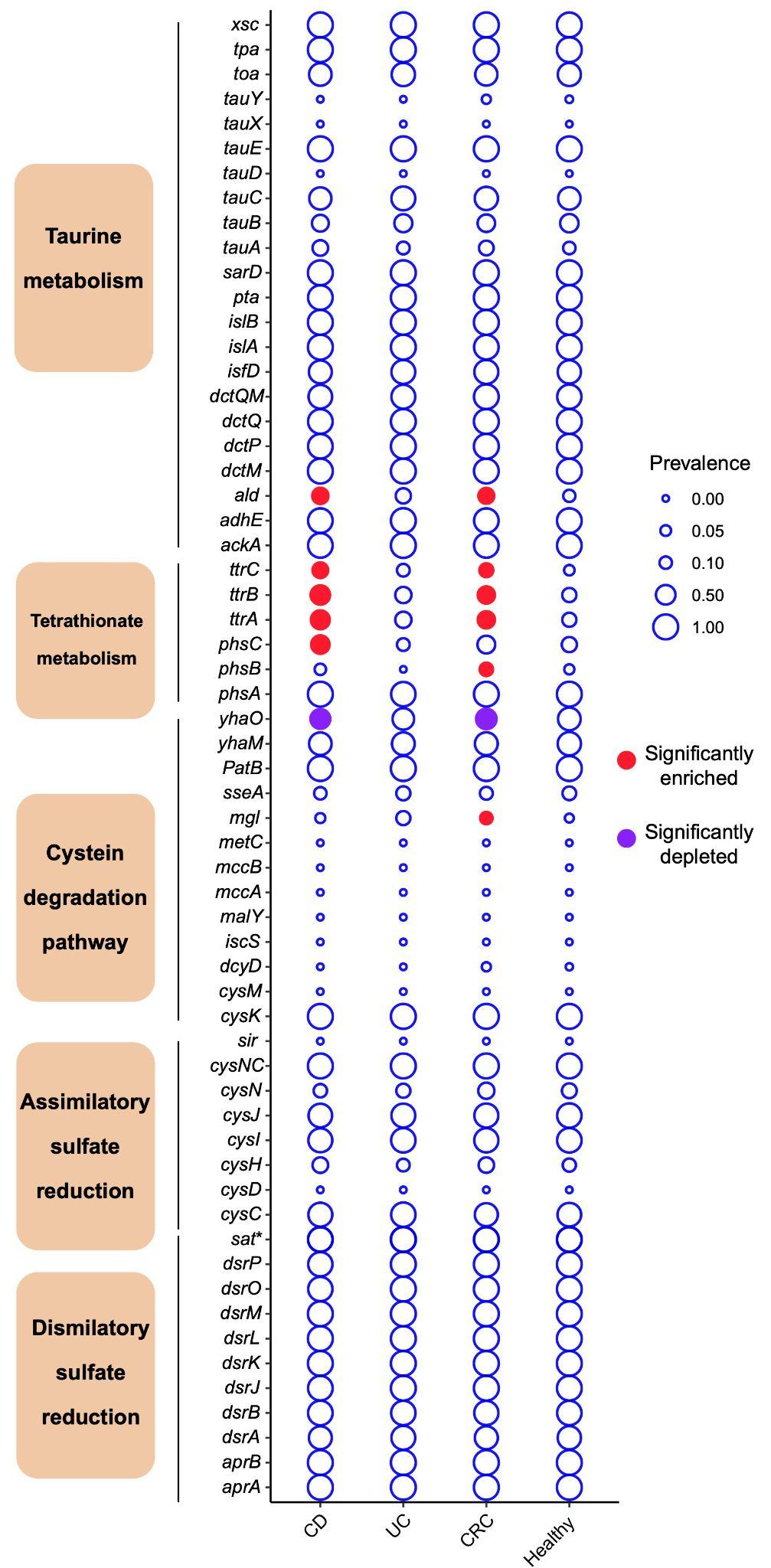

**Supplementary Fig. 16. Prevalence of sulfur metabolic genes across all *Desulfovibrio* genomes.** Sulfur metabolic genes were identified by an hmm search of a curated list of sulfur metabolic genes. The genes that are significantly enriched (red) or depleted (purple) in IBD or CRC are highlighted (Fisher's exact test, adjusted p-value <0.25).*: *sat* is involved in dissimilatory and assimilatory sulfate reduction (KEGG modules M00596 and M00176).

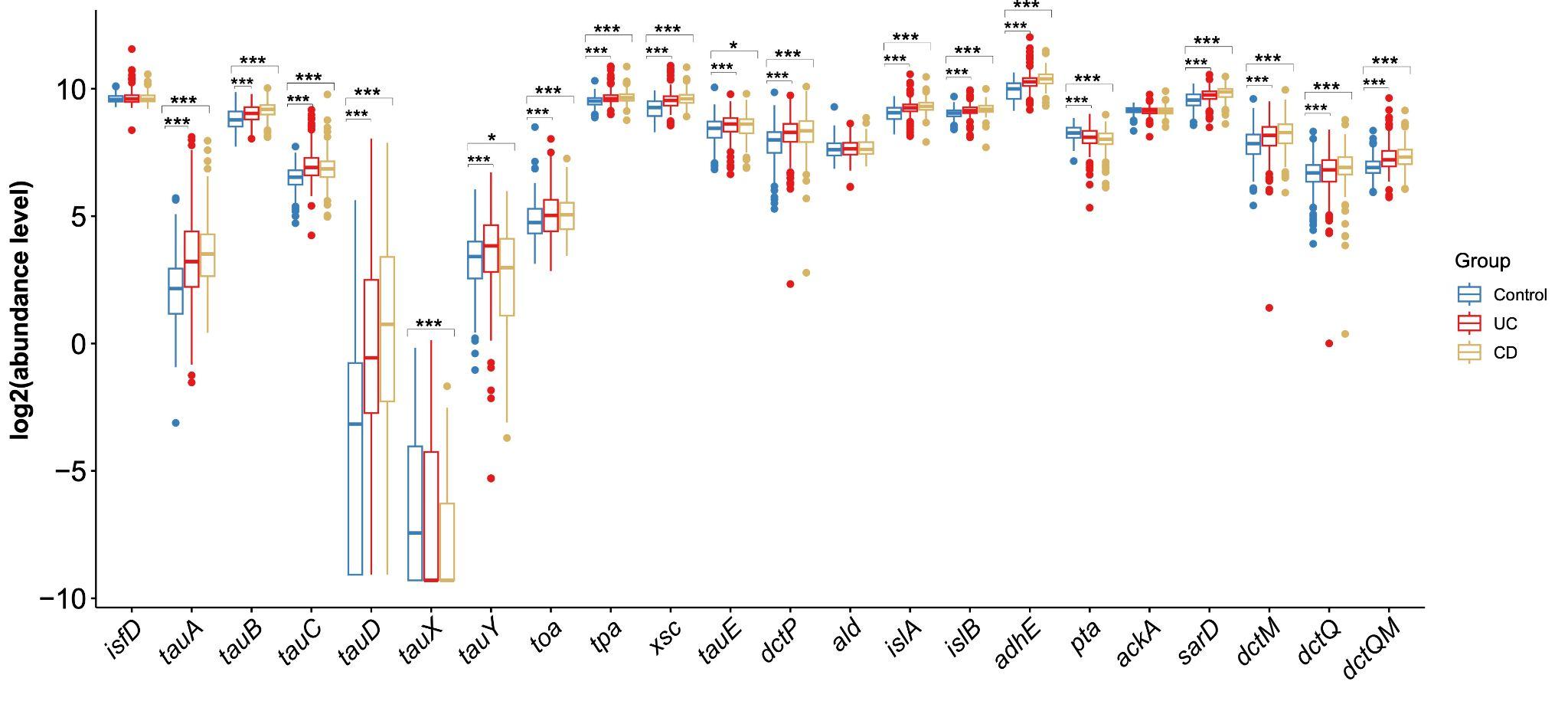

**Supplementary Fig. 17. Most genes involved in taurine metabolism were differentially abundant in IBD compared to the control group at the microbiome community level.**  Differential abundance analysis was performed on all genes involved in the taurine metabolism (Wilcoxon test). Stars represent different significance level: * p < 0.05; ** p<0.01; *** p<0.001.

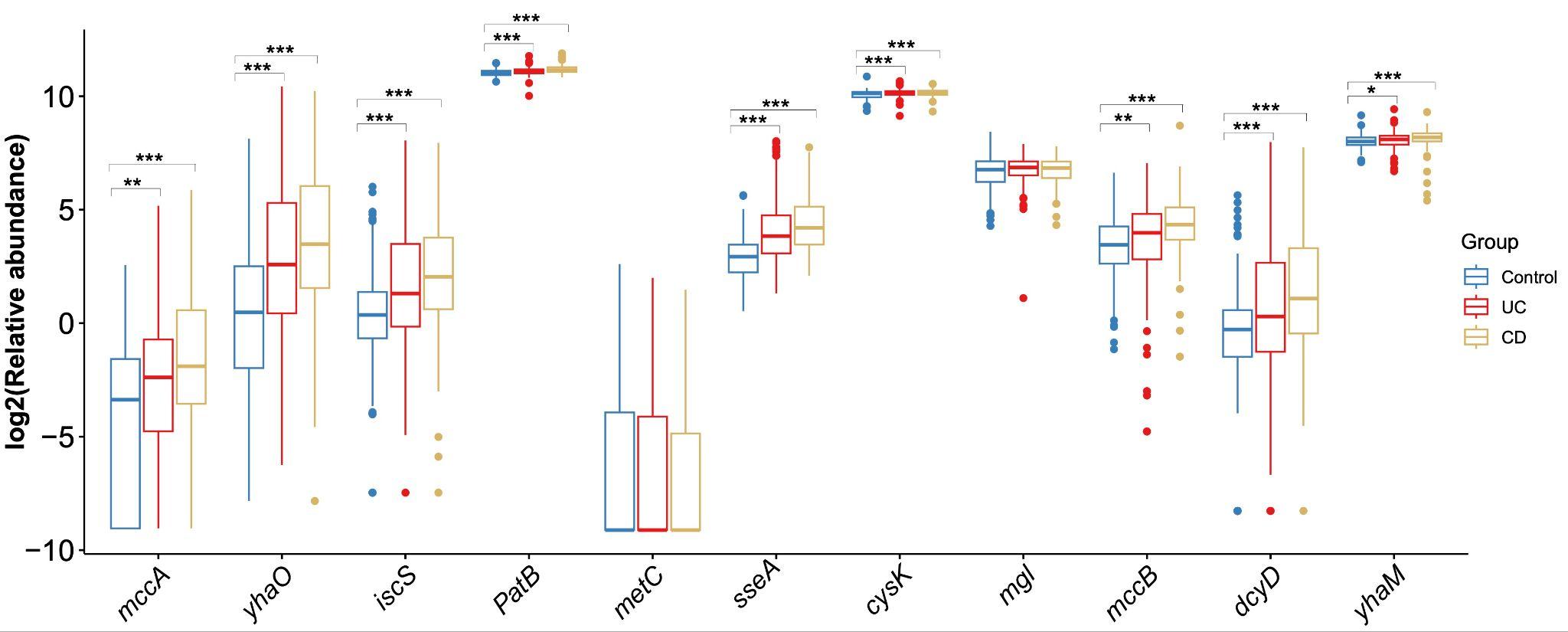

**Supplementary Fig. 18. Most genes involved in cysteine degradation were differentially abundant in IBD compared to the control group at the microbiome community level.** Differential abundance analysis was performed on all genes involved in cysteine degradation (Wilcoxon test). Stars represent different significance level: * p < 0.05; ** p<0.01; *** p<0.001.

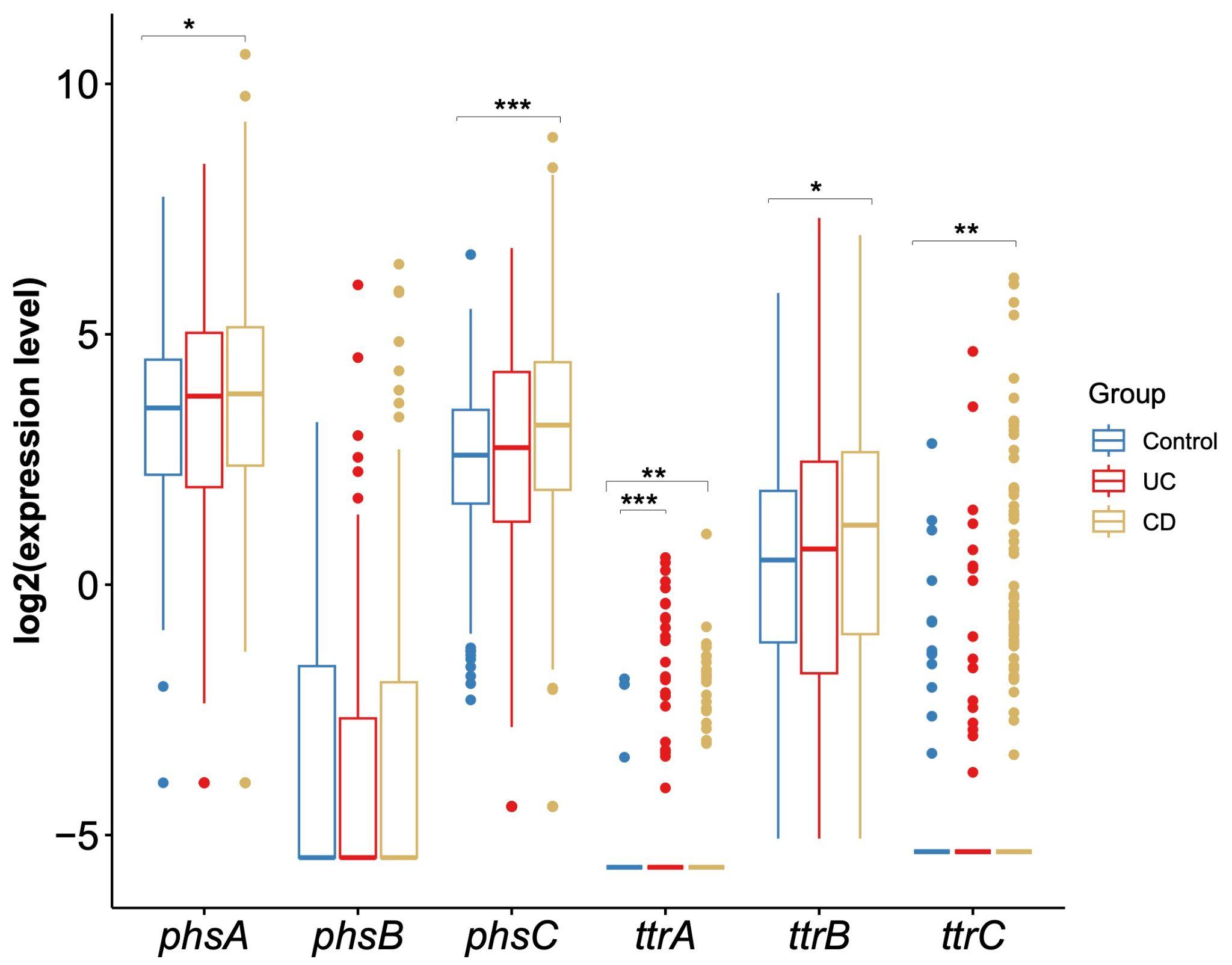

**Supplementary Fig. 19. Metatranscriptomic data indicated that most genes involved in tetrathionate metabolism were significantly upregulated in CD compared to non-IBD controls.** Differential expression analysis was performed on all genes involved in the tetrathionate metabolism pathway (n = 6, Wilcoxon test). Stars represent different significance level: * p < 0.05; ** p<0.01; *** p<0.001.

### **Supplementary Tables**

**Supplementary Table 1. Statistics and host information for *Desulfovibrio* genome assemblies.**

**Supplementary Table 2. Metagenomic datasets used for new *Desulfovibrio* MAG identification.**

**Supplementary Table 3. *Desulfovibrio* prevalence in cohorts used for de novo MAG discovery.**

**Supplementary Table 4. Genomes reassigned to other genera within the Desulfovibrionaceae family using GTDB-tk.**

**Supplementary Table 5. Detailed information on the origin, availability and assembly statistics for the 24 human *Desulfovibrio* isolates.**

**Supplementary Table 6. Overview of all studies used to compile the *Desulfovibrio* database.**

**Supplementary Table 7. *Desulfovibrio sp900319575* genes that were enriched in the Netherlands and East Asia respectively.**

**Supplementary Table 8. Clusters of Orthologous Genes (COGs) significantly enriched in Crohn's disease (CD) vs healthy individuals (Fisher's exact test, adjusted p-value <0.25).**

**Supplementary Table 9. Clusters of Orthologous Genes (COGs) significantly enriched in colorectal cancer (CRC) vs healthy individuals (Fisher's exact test, adjusted p-value <0.25).**

**Supplementary Table 10. Hmm files for the curated microbial sulfur metabolism gene database.**

**Supplementary Table 11. MAGs identified in a deeply sequenced IBD cohort (FedericiS_2022).**
